# Coding advantage of grid cell orientation under noisy conditions

**DOI:** 10.1101/507061

**Authors:** Dong Chen, Kai-Jia Sun, Liang Wang, Zhanjun Zhang, Ying-Cheng Lai, Nikolai Axmacher, Wen-Xu Wang

## Abstract

Grid cells constitute a crucial component of the “GPS” in the mammalian brain. Recent experiments revealed that grid cell activity is anchored to environmental boundaries. More specifically, these results revealed a slight yet consistent offset of 8 degrees relative to boundaries of a square environment. The causes and possible functional roles of this orientation are still unclear. Here we propose that this phenomenon maximizes the spatial information conveyed by grid cells. Computer simulations of the grid cell network reveal that the universal grid orientation at 8 degrees optimizes spatial coding specifically in the presence of noise. Our model also predicts the minimum number of grid cells in each module. In addition, analytical results and a dynamical reinforcement learning model reveal the mechanism underlying the noise-induced orientation preference at 8 degrees. Together, these results suggest that the experimentally observed common orientation of grid cells serves to maximize spatial information in the presence of noise.

**Author summary:** Spatial navigation depends on several specialized cell types including place and grid cells. Grid cells have multiple firing fields that are arranged in a regular hexagonal pattern. The axes of this pattern are anchored to environmental boundaries at a universal angle of 8°. Here, we combine computer simulations of the grid cell network with analytical derivations and a reinforcement learning model to explain the functional relevance of this universal grid cell orientation. We show that spatial information provided by grid cells is maximized at the experimentally observed grid orientation within a broad parameter range. This relationship occurs only in the presence of noise. The model allows for several experimentally testable predictions including the number of grid cells.

## Introduction

The past decade has witnessed substantial progress in our understanding of the internal positioning system (“GPS”) in the mammalian brain. Specifically, place cells in the CA1 region of the hippocampus convey unique information about spatial position [1-2], while grid cells implement a universal coordinate system that provides information about spatial distances [3]. In addition, a number of other functionally dedicated cells were found to play a role in creating the neural representation of space, such as border cells [4-5], head direction cells [6], and speed cells [7]. Among all the known types of cells that constitute the brain’s GPS, grid cells have received particular attention due to their abundant number, their determining role in generating an inner metric for navigation [8-10], and their close relationship to Alzheimer’s disease [11-12].

A large number of experiments in animals and humans have demonstrated that grid cells exhibit multiple hexagonally arranged firing fields that tile the entire space (Fig 1A). Because of the efficient geometrical organization of the firing fields, activity of a small number of grid cells is sufficient to provide a nearly full coverage [3, 13-14]. In addition to the hexagonal geometry of the firing patterns, experiments revealed two additional striking features: First, grid cells can be assigned to 4-5 discrete modules that are each defined by a common spatial scale and orientation but different phases [3, 15] (Fig 1B). Second, the spatial scale increases geometrically across modules [15], with a fixed geometrical ratio of around 1.4-1.7. Several computational models have been developed to explain the firing patterns of grid cells. Attractor network and oscillatory interference models were proposed to explain the hexagonal geometry [16-17], and models considering the maximization of spatial information were introduced to understand the geometrical ratio of spatial scales [18-20]. These studies demonstrated that spatial locations can be decoded from the population activity of grid cells from each modules, and that the hexagonal firing patterns and geometric scale ratios maximize spatial information content. Overall, these results offer a possible explanation for the evolutionary significance of the functional properties of grid cells.

**Fig 1.**
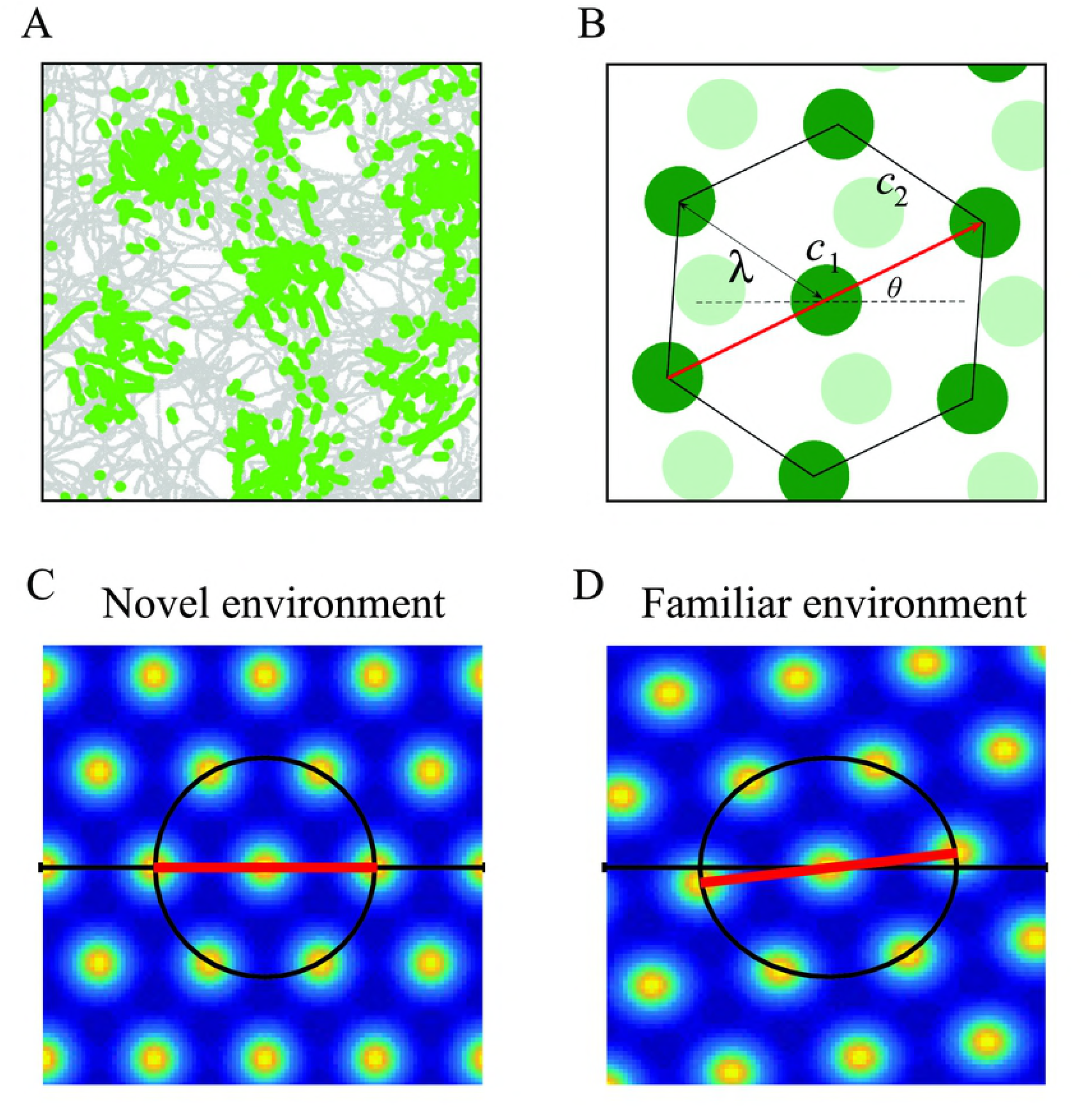
Spatial firing fields of grid cells. (**A**) Typical firing fields of a single grid cell in an experiment, where the firing locations are marked in green dots superimposed on the trajectory (gray) of a rat (data from http://www.ntnu.edu/kavli/research/grid-cell-data). (**B**) Schematic illustration of the hexagonal firing pattern (dark green color), which is defined by the orientation angle *θ* of the grid (red arrow), the module scale *λ* (black arrow), and the spatial phase ***c***_1_. The firing field of another grid cell (light green) within the same module shares the common orientation and scale, but has a different phase ***c***_2_. (**C**) Simulated firing fields with *θ* = 0 degrees and an ideal symmetric hexagonal array (black circle) in a square environment. (**D**) Simulated firing fields matching experimental results. Orientation *θ* deviates from 0 degrees and there is an elliptic distortion of the firing pattern.

Although the hexagonal geometry and the modular structure generated by grid cells have been reasonably well understood, an equally fundamental issue that remains elusive is the orientation of the hexagons. Specifically, are the hexagonal grids anchored to an external reference frame – and if so, which mechanisms account for anchoring, and which purpose does it serve? Intuitively, since the hexagonal scale can vary from module to module and may adapt to the environment [21-22], one might speculate that the orientation of different modules would be different. However, recent experiments revealed the striking phenomenon that, in a square environment with closed boundaries, a specific orientation of the hexagonal firing grids emerges [23-25], which is universal across different grid cells, modules and subjects. The common orientation anchored to the square borders may be associated with evolutionary significance and stem from natural selection. In fact, in three independent experiments, the measured orientation angles were found to be quite close: they are clustered at 7.4 degrees [23], 8.8 degrees [24] and 8.2 degrees [25], respectively. This specific orientation is accompanied by a distortion of the hexagonal grid (e.g., comparing Fig 1C with Fig 1D), which was attributed to shearing forces resulting from an interaction with the domain borders [23]. Yet, an explicit explanation of the biological significance of the specific orientation is lacking, and the mechanism underlying the universal orientation at around 8 degrees is unknown [26-27]. Addressing these questions will likely yield important insights into how grid cells interact with border cells in order to adapt to the environment [28-29], and more generally will advance our understanding of the function of grid cells for spatial coding [30].

In this paper, we propose a spatial coding scheme based on the population activities of grid cells pertaining to different modules, and hypothesize that the universal and stable orientation at 8 degrees maximizes the amount of spatial information conveyed by grid cells. Combining an encoding model, theoretical analyses and a reinforcement learning model, we provide converging evidence that an orientation of 8 degrees maximizes the amount of spatial information conveyed by grid cells in the presence of noise. Additionally, our computer model allows for empirical testable predictions on the minimal number of grid cells required for an optimal encoding scheme.

## Results

The hexagonal firing pattern of a single grid cell, as exemplified by each of the seven dark green circles in Fig 1B, can be defined by the orientation, the scale, and the spatial phase of the firing field. Experiments with rats revealed that grid cells are organized into 4 or 5 functional modules, within which the firing fields of the cells share the same scale and orientation but differ in spatial phases, as shown in Fig 1B. The scale increases geometrically between the discrete modules. In general, a single module is insufficient to provide unambiguous information about spatial locations but the population activities of grid cells from all modules can represent spatial locations at a high resolution [20]. Previous models suggested that, in a circular environment, the hexagonal array and geometric progression of scale is optimal with respect to efficient spatial coding [19]. In contrast, the underlying mechanisms and functional roles of a grid cell’s orientation has not yet been fully understood. Interestingly, recent experiments revealed a universal orientation of the firing fields of about 8 degrees with respect to the borders of a square environment [23-25]. In addition, displacement of borders induce changes in orientation, and these are accompanied by distortions of the hexagonal firing grid. Both effects can be conceived of as resulting from shearing forces from the borders [23]. In order to understand the role of grid orientation quantitatively, we develop a spatial encoding model without any adjustable parameters.

We hypothesize that the universal orientation optimize spatial encoding. In particular, we note that the ubiquity of grid cells in mammals [31-34] suggests certain evolutionary advantages of their properties. Likewise, the specific orientation with respect to borders may also be evolutionarily significant for optimizing the information content of spatial representation. To reveal quantitatively how the population activities of grid cells across different modules support behavior in the positioning and self-localization tasks, we study spatial encoding via population activities.

### Spatial representation of the grid cells system

The spatial representation via grid cells is modeled as follows. Assume there are 4 modules with spatial scales *λ*(*k*), for *k* =1,2,3,4, and a fixed geometrical ratio *r* of the scales between two adjacent modules: *λ*(2)/*λ*(1) = *λ*(3)/*λ*(2) = *λ*(4)/*λ*(3) = *r* > 1. Each module consists of a regular, topographically ordered population of *M* cells [20, 35]. The firing fields of *M* cells tile periodically in an environment. Since we still lack exact experimental evidence for the number of cells *M*, we will study the effect of varying *M* on the spatial information and the orientation of grid patterns. For an arbitrary grid cell *i* from a module *k*, let *θ* be the smallest angle between one of the six symmetric axes of the hexagonal firing pattern and four borders. The ideal spatial firing rate 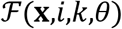 at an arbitrary location **x** can be conveniently represented by a bell-shaped, spatially periodic function on a hexagonal lattice [20] (see Methods). The spatial distribution of 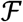 is illustrated in Fig 1C. This establishes a mathematical description of the spatial firing fields of all grid cells and allows investigating how spatial locations in an environment are encoded based on their firing patterns. Because the firing fields of a single cell repeat across environments, the activity of that cell does not provide an unambiguous representation of self-location. However, the conjunction of cells across 4 modules can be unambiguous [20].

Based on the firing rate 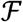, the firing intensity *I*(**x**,*i,k,θ*) of cell *i* at a given location **x** can be measured within a time interval *τ*:

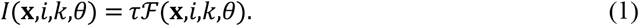

Because of several intrinsic sources of neuronal noise, such as stochastic channel openings, thermal noise and noise due to synaptic background activity [36-37], the firing intensity *I*(**x**,*i,k,θ*) is not deterministic but stochastic. We assume spiking to follow a Poisson distribution with expected value *τ* 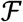 [14, 18-20] (see Methods). Because stochastic fluctuations of time courses that follow a Poisson distribution cancel out with increaseing durations, for a longer time interval *τ* the signal to noise ratio (SNR) will be larger. We use the maximum value of firing intensity *τf*_max_ to measure the strength of the signal and the standard deviation 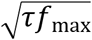 to quantify noise. Thus, the effect of noise can be evaluated by the ratio of noise to signal, i.e., the inverse of 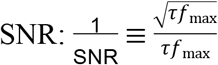. Since *f*_max_ is unchanged, the effect of noise is reduced as *τ* increases. Below, we will use 1/SNR to characterize the effect of noise and its inverse correlation with *τ*. The larger the value of 1/SNR, the stronger the effect of noise.

### Spatial encoding and evaluation

Our proposed spatial encoding scheme based on the population activities of grid cells across different modules includes intra-module encoding and inter-module encoding, where the former is associated with the assumption that grid cells in the same module compete to represent spatial locations, and the latter stems from the assumption that cells in different modules complement each other for spatial representation. Although our hypothesis lacks immediate and strong experimental supports, some evidence in the literature indicates this simple encoding scheme is consistent with the empirical properties of grid cells. Specifically, the intra-module encoding scheme implements a “winner-take-all” (WTA) rule: a location is represented and encoded by the cell with the highest firing intensity *I* at the location. WTA rules are typically implemented by lateral inhibition [38] consistent with the experimentally observed coupling of grid cells via inhibitory interneurons [39] that constitute the essential element of attractor models of hexagonal firing patterns of grid cells [40]. The WTA rule can be formulated as:

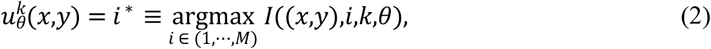

where *k* =1,2,3,4, correspond to 4 modules, and *i*\* is the index of the cell that dominates (encodes) location (*x,y*) in module *k*. Fig 2A illustrates the WTA in the absence of noise. As the result of WTA, each grid cell has a set of periodically repeated dominant sites with a hexagonal shape in a module, and the center of each dominant site coincides with that of the original firing fields of the grid cell (Fig 2A). At any given location, there are four elements 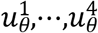 that constitute a vector **u**_*θ*_ for encoding this location, as illustrated in Fig 2B. In general, it can be speculated that more cells and modules enable a more precise encoding of locations in terms of less duplication of vector **u**_*θ*_, such that each location can be more explicitly distinguished from others based on the coding scheme. However, regarding the biological cost of creating neurons and supporting their metabolism, there must be a tradeoff between the number *M* of grid cells and the efficiency of spatial encoding.

**Fig 2.**
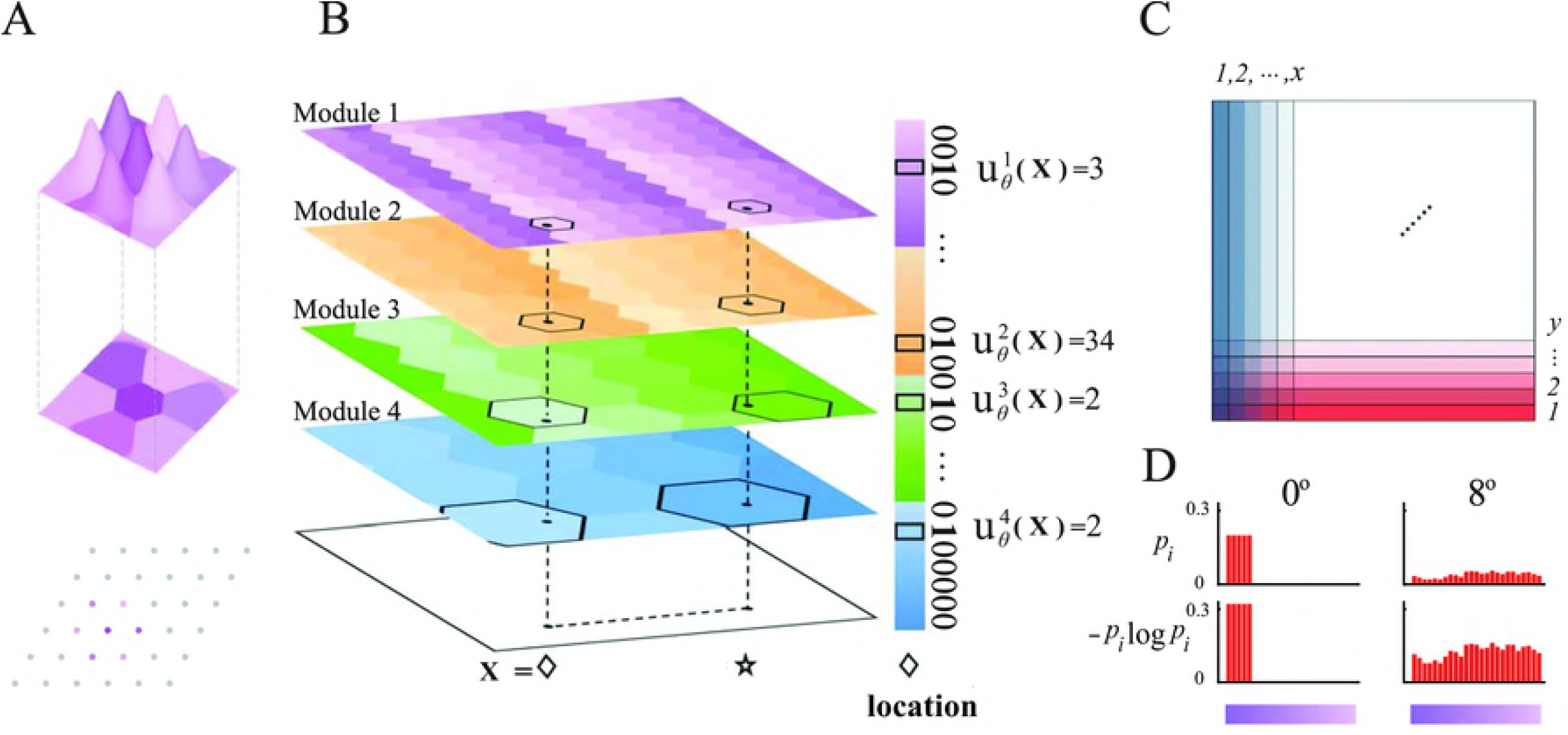
Encoding scheme of multiple modules and spatial information entropy. (**A**) Top: Example of a winner-takes-all (WTA) induced dominant firing site (encoded site) with a hexagonal shape in the center, resulting from competition between the original firing fields among neighboring grid cells according to the WTA rule. Below: the centers of the dominant cites encoded by 6^2^ grid cells (*M* = 6^2^) in a spatial unit with a rhombus shape in a module. The spatial unit with the encoded sites iterates periodically in the space. (**B**) Dominant sites encoded by grid cells with different colors in modules 1 to 4. Every dominant firing site has a hexagonal shape. The encoded sites in a spatial unit are arranged periodically in every module. Every location is encoded by four sites (highlighted in each module) from four modules, respectively, e.g., the two locations marked by diamond and star. The four encoded sites of location ◊ constitute a spatial vector *u_θ_*(◊), where each of the four elements denotes the index of the cell engaged in encoding the site pertaining to location ◊ in a module. (**C**) The whole space is divided into 2*N* belts, where the *N* blue and red belts are parallel to the two sets of borders, respectively. Every belt consists of *N* boxes with unit scale. (**D**) Illustration of the occurrence probability *p_i_* and the spatial information entropy – *p_i_*log*p_i_* of different cells in the two belts with *θ* = 0 and 8 degrees, respectively. The color bar represents the index of different cells.

To evaluate the capacity of this spatial encoding scheme in a closed environment, we resort to the concept of information entropy. Let us consider two arbitrary spatial vectors **u**_*θ*_(**x_1_**) and **u_*θ*_**(**x_2_**). If **u**_*θ*_(**x_1_**) = **u**_*θ*_(**x_2_**), locations **x_1_** and **x_2_** will be considered the same and cannot be differentiated. In a space, a higher number of indistinguishable locations is associated with a poorer spatial representation and a lower encoding capacity. Information entropy is a parsimonious metric for accessing encoding capacity in terms of measuring the amount of effective information. Note that spatial information encoded from different modules is considered to be independent, so that the different modules complement each other. Thus, we calculate information entropy of each module separately.

Moreover, we divide the square domain along borders into a number of belt regions of unit width, as shown in Fig 2C. We obtain two groups of belt regions, parallel to the two groups of borders of the square domain, respectively. Our hypothesis of the division is that a two-dimensional domain can be reduced to two orthogonal directions (one dimension each) by simply anchoring to borders. The encoding of two dimensions can complement each other to represent the whole space. In addition, border cells may help grid cells encode locations close to borders. Taken together, if a spatial encoding scheme generated by grid cells is able to identify locations within each belt, all locations in the domain can be explicitly distinguished and an efficient spatial encoding is established.

The capacity of spatial encoding within a belt relies on the spatial distribution of sites 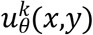 encoded by grid cells. Let us consider two extreme scenarios. On the one hand, if within a belt, every location is represented (encoded) by the same cell, i.e., there is a single value of *u*, the spatial representation is maximally poor and any two locations cannot be distinguished. In this extreme scenario, the capacity of spatial encoding reaches the lower bound according to information theory. On the other hand, if every location is uniquely encoded by a cell (the value of *u* is completely different across all different locations), the capacity of encoding locations will be maximal. The encoding capacity in the scenarios can be effectively measured by information entropy within a belt. To be concrete, we denote two groups of belts by index *x* and *y*, respectively, as shown in Fig 2C. Here each cell is akin to a random variable and its frequency of appearance in a belt corresponds to the probability of a random variable, denoted by a conditional probability *p*(*i*|*θ,k,x*) of cell *i* from module *k* in a given belt *x*:

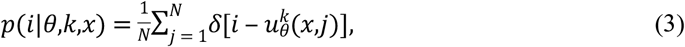

where *δ* is the Dirac function with *δ*(*l*_1_ – *l*_2_) = 1 if *l*_1_ = *l*_2_ and *δ*(*l*_1_ – *l*_2_) = 0 otherwise, *j* is the index of the unit box within belt *x* and *N* is the total number of boxes in belt *x*. The right hand side of formula (3) denotes the probability of finding boxes encoded by cell *i*. Analogously, the occurrence probability *p*(*i|θ,k,y*) in the other group of belts *y* can be defined by substituting 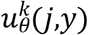 for 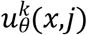.

The spatial information entropy *E* of a belt *x* is defined as:

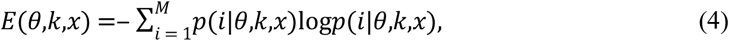

where *i* is the cell index and *M* is the number of cells in each module. *E* of a belt *y* is defined in the same way. Examples of *p* and *E* of two belts with *θ* = 0 degrees and 8 degrees are illustrated in Fig 2D. *E*(*θ,k*) of module *k* is the summation of the average *E* of belts along *x* and *y* directions, respectively:

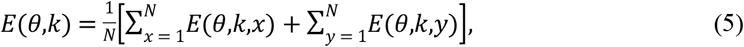

where *N* is the number of belts along each direction. Finally, the information entropy *E* of a domain is the average over 4 modules:

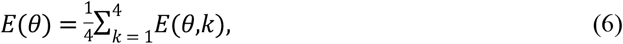

which depends on the orientation *θ*.

We are able to explore how *E*(*θ*) is affected by orientation *θ*, taking into account different internal and external conditions, such as the number of cells *M* in each module, geometric ratio *r*, effect of noise 1/SNR, elliptic distortion *ε* of hexagonal grid, and size *N* of the square environment. Because the hexagonal grid has a rotational symmetry, the orientation, defined as the minimal angle of the grid from any border of a square space, can be conveniently confined to the range [0,15] degrees [*23*].

### Numerical results of optimal orientation angle

Our first goal is to numerically identify the optimal orientation angle *θ*_opt_ that maximizes information entropy *E* with respect to different internal and external conditions.

Fig 3A shows examples of *E* as a function of *θ* for two values of *M*, a typical geometrical ratio *r* and size of space in the ideal situation without noise and elliptic distortion. We see that a global peak of *E* arises at *θ* = 8 degrees and a local valley at *θ* = 11 degrees. A more comprehensive assessment of optimal *θ*_opt_ as a function of both *r* and *M* is shown in Fig 3B. We see that *θ*_opt_ is insensitive to *r* within the range 1.4-1.7, but strongly depend on *M*: only in the region *M* ∈ [7^2^,13^2^], *θ*_opt_ = 8 degrees. Outside of this small region, *θ*_opt_ significantly deviates from 8 degrees. Fig 3C shows the average standard deviation *σ* of *E* among belts along *x* and *y* directions at *θ* = *θ*_opt_. We see that *σ* is relatively negligible, indicating that *E* of belts along each direction is generally stable, regardless of *θ*_opt_. (Results for other typical environment sizes can be seen in S1A-B and F-G Fig.)

**Fig 3.**
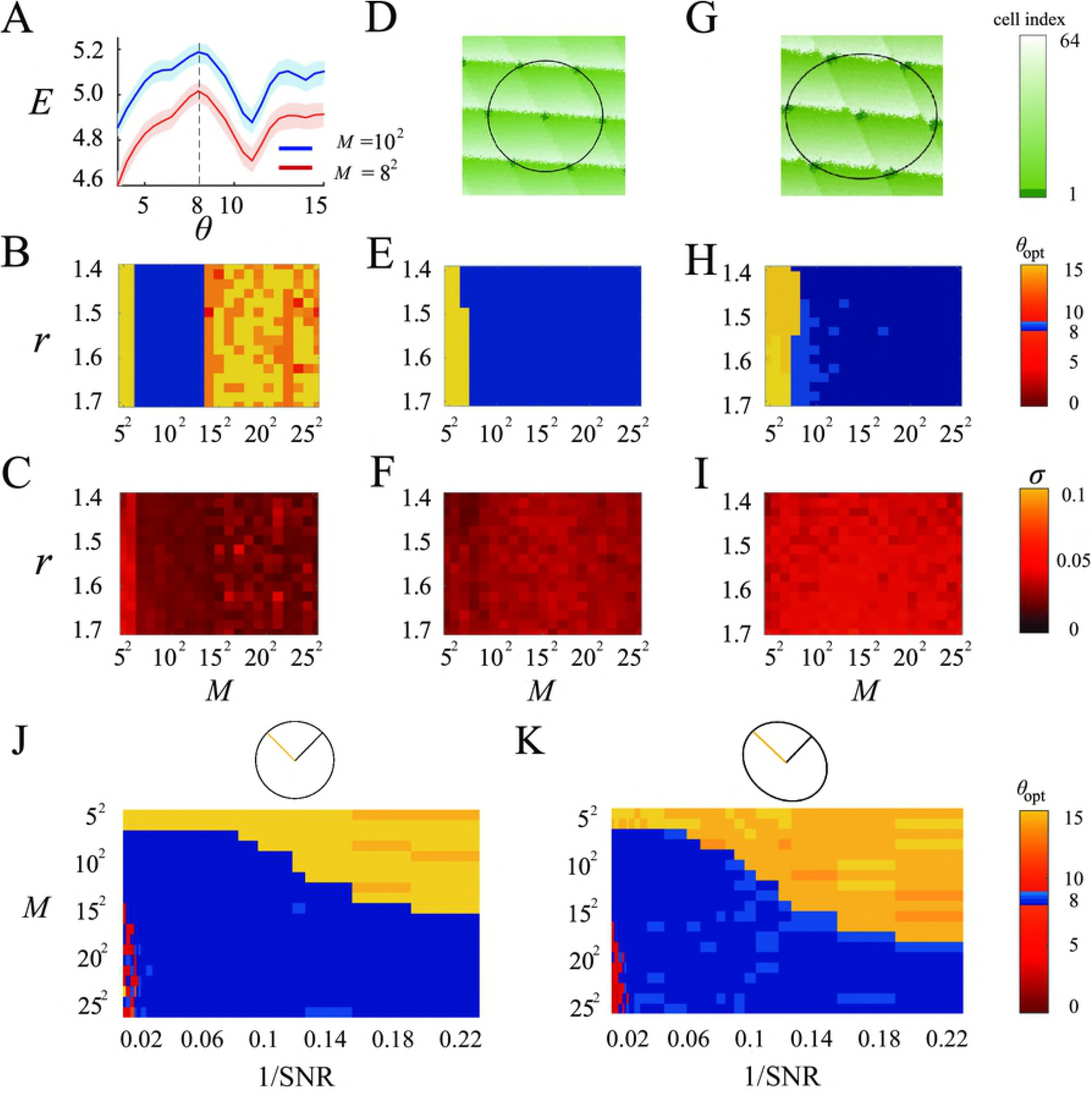
Optimal orientation associated with highest spatial information entropy. (**A**) Spatial information entropy *E* of a square space as a function of orientation *θ* for *M* = 8^2^ and *M* = 10^2^ in an ideal situation without noise and elliptic distortion. Geometrical ratio *r* = 1.44. The shadow areas represent standard deviation among different belts. (**B**) Optimal orientation *θ*_opt_ associated with highest *E* as a function of both *r* and *M* in an ideal situation. *θ*_opt_ occurs at 8 degrees in the range 7^2^ ≤ *M* ≤ 13^2^ and is largely insensitive to *r*. (**C**) Standard deviation *σ* of *E* among different belts at *θ*_opt_ as a function of both *r* and *M* in the ideal situation. (**D**) Illustration of spatial encoding pattern *u* based on the WTA rule in a module in the presence of noise without elliptic distortion. (**E**)-(**F**) *θ*_opt_ (**E**) and standard deviation *σ*(*E*) among different belts (**F**) as a function of both *r* and *M* in the presence of noise without elliptic distortion. (**G**) Illustration of spatial encoding pattern *u* in the presence of both noise and elliptic distortion. (**H**)-(**I**) *θ*_opt_ (**H**) and *σ* of *E* among different belts (**I**) as a function of both *r* and *M* in the presence of both noise and elliptic distortion. (**J**)-(**K**) *θ*_opt_ as a function of both the effect of noise 1/SNR and grid numbers *M* in the presence of noise without elliptic distortion (**J**) and in the presence of both noise and elliptic distortion (**K**). In (**E**) and (**H**), *θ*_opt_ = 8 degrees when *M* > 8^2^, largely independent of *r*. In (**D**) to (**I**), the effect of noise 1/SNR = 0.08. In (**G**) to (**I**) and (**K**), elliptic distortion intensity *ε* = 1.17 as observed in experiments (23). The color bar of (**B**), (**E**) and (**H**) represents *θ*_opt_. The color bar of (**C**), (**F**) and (**I**) represents standard deviation *σ*. In (**J**)-(**K**), the geometrical ratio *r* = 1.5 and in all the results, the environmental size is 1.5 × 1.5m^2^, *f*_max_ = 200Hz and *τ* =0.8s. The duration time *τ* corresponding to 1/SNR < 0.22 is *τ* > 100ms.

It is worth noting that *M* represents the actual number of grid cells in each module. Our simulations suggest that *ME* [7^2^,13^2^]. Notably, these numbers are independent from any assumptions on the empirical values of cell numbers *M*; the exclusive criterion to obtain *M* is the maximization of information entropy at orientations of about 8 degrees. Interestingly, the range of *M* ∈ [7^2^,13^2^] – i.e., cell numbers between about 50 and 170 per module – is substantially smaller than previous speculations about the numbers of grid cells, which assume about 70,000 cells in layer II of EC [42] of which are about 10% grid cells [41], resulting in 7,000 grid cells across 5-7 modules [14, 19]. This suggests that either the number of grid cells is smaller than previously estaimated, or that some grid cells have other functions than the optimizaton of spatial information.

Next, we considered the effect of noise embedded in the firing intensity on *θ*_opt_. Fig 3D shows the spatial encoding pattern of *u*(*x,y*) blurred by noise in a module. To our surprise, spatial noise significantly enlarges the parameter region in which *θ*_opt_ = 8 degrees, as shown in Fig 3E as an example. Below, we will theortically assess the mechanisms driving this effect of noise on *θ*_opt_. Fig 3F shows that *E* of belts are still stable, irrespective of the effect of noise and other parameters. The stability of *E* among belts facilitates our theoretical analyses by allowing us to arbitrarily choose two representative belts from the two directions, as explained in detail below. (Results for other two typical environment sizes can be seen in S1C-D and H-I Fig.)

In addition to noise, we incorporated elliptic distortion to fully capture real hexagonal grids as described experimentally (see Methods for the definition of the intensity of elliptic distortion *ε*). Fig 3G depicts the spatial encoding pattern of *u*(*x,y*) in a module under the influence of both noise and elliptic distortion. Simulation results show that the effect of distortion is negligible, because that both *E* and *σ* take very similar values as in the condition without distortion (compare Fig 3E/F to Fig 3H/I). (Results for other two typical values of distortion intensity *ε* can be seen in S2 Fig.)

Next we study the effect of noise 1/SNR on optimal orientation *θ*_opt_. Since *E* is insensitive to *r*, we ignore *r* and take into account both noise effect 1/SNR and cell numbers *M*. Fig 3J shows *θ*_opt_ as a function of 1/SNR and *M*. Strikingly, we find that the parameter map of *θ*_opt_ is clearly dominated by an orientation of 8 degrees in a large range of 1/SNR from 1/SNR→0 to 1/SNR = 0.22. Specifically, there exists a lower bound of *M* versus 1/SNR, i.e., when *M* exceeds the bound, *θ*_opt_ always occurs at 8 degrees. The lower bound of *M* increases slightly from 7^2^ to 15^2^ as 1/SNR is increased from 0 to 0.22. These results demonstrate that noise indeed provides strong support for the emergence of *θ*_opt_ at 8 degrees in a wide range of noise levels and the number *M* of grid cells. Moreover, the prediction of the lower bound of *M* is relevant for the interpretation of experimental results, since examining *M* through experiments is still challenging at present. Fig 3K shows *θ*_opt_ as a function of both 1/SNR and *M* with elliptic distortion. The results resemble those in Fig 3J without distortion, indicating the negligible effect of elliptic distortion on *θ*_opt_ compared to noise. (See also S1E, J Fig and S2E, J Fig.) Taken together, simulation results demonstrate that in an ideal situation without noise and distortion, *θ*_opt_ occurs at 8 degrees only in a relatively small region of *M*. In contrast, in the presence of noise and elliptic distortion, an orientation of 8 degrees dominates *θ*_opt_ as long as *M* exceeds some lower bound – specifically, noise plays a major role in the prominence of 8 degrees, while the effect of elliptic distortion is negligible. As an alternative to information entropy, the capacity of this encoding scheme can be characterized by means of Hamming distance among the location vectors (see S1 Methods and S3 Fig for details). Specifically, if the Hamming distance between the vectors of two locations is larger, the two locations can be more explicitly discriminated. Thus, the average Hamming distance among all pairs of location vectors in an environment can capture the ambiguity of encoding the environment, and the larger Hamming distance, the lower ambiguity. There is an intrinsic correlation between information entropy and the Hamming distance, since both of them evalutate spatial ambiguity. In general, the higher information entropy, the larger the average Hamming distance among locations. However, the Hamming distance is not linearly proportional to information entropy. The Hamming distance yields the same result of optimal orientation as that based on the information entropy, providing additional evidence to validate our encoding scheme and the effect of noise on optimal orientation (see S4 Fig).

### Correlation between information entropy and the number of cells engaged in encoding

The purpose of studying the relationship between *E* and the number of cells *n_c_* engaged in encoding a belt should eventually enable for an analytical assessesment of *θ*_opt_. Note that *n_c_* is equivalent to random variables in the standard definition of information entropy. According to Shannon’s theory, the occurrence probability of a variable is inversely correlated with the amount of effective information encoded by this variable. In other words, a higher number of random variables usually results in a higher information entropy. This is because in general, adding more variables decreases the average probability of each variable, such that their encoding capacity is enhanced. Thus, we speculate that *E* of a belt is positively correlated with *n_c_* of the belt. As shown in S5A Fig, indeed an approximately positive linear correlation arises between *n_c_* and *E*, and the correlation is largely independent of the exact values of *M*. We also test how the correlation is affected by noise and elliptic distortion. The results shown in S5B Fig indicate that the correlation is robust against noise, reflected by an only slight decrease as the effect of noise 1/SNR increases. The elliptic distortion has negligible effect on the correlation as well (S5B Fig). The high and robust correlation between *E* and *n_c_* allows us to substitute *n_c_* for the more complicated *E* to obtain analytical results of *θ*_opt_.

### Geometrical analysis of optimal orientation in a noise- and distortion-free situation

We develop a heuristic analysis to theoretically predict *θ*_opt_ = 8 degrees, for *M* ∈ [7^2^,13^2^] in the absence of noise and elliptic distortion. Our theoretical approach is basically a geometrical analysis relying on the strong correlation between *E* and *n_c_* and the strong stability of *E* among different belts, where the latter allows us to arbitrarily select two belts from *x* and *y* directions that represent an entire two-dimensional square. For simplicity, we use two orthogonal and sufficiently long lines without widths to stand for the two belts, as shown in Fig 4A. Each line will traverse a number of hexagonal sites *u* encoded by grid cells. Note that the number of cells *n_c_* engaged in encoding along a line can be estimated by the mean repeating distance between two nearest locations encoded by a given cell. Specifically, every location within the repeating distance along a line will be encoded by a number of different grid cells. Thus, a longer repeating distance requires a larger number of grid cells *n_c_* to encode every location within the repeating distance. However, the spatial distribution of *u* along a line for a given *θ* is rather complex, such that it is difficult to analytically assess the mean repeating distance. Therefore, we reduce the mean repeating distance to the minimum repeating distance among all encoded sites along a line to approximate *n_c_*. Our analytical results validate this approximation. Let us denote the minimum repeating distance of all hexagonal sites *u* along the two lines by *T*(*θ*) and 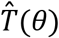 for an arbitrary *θ*. This yields the relation

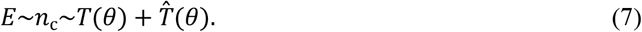

We numerically found high correlations between *E* and 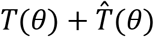, and between *n_c_* and 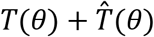, as shown in S6 Fig. We find that within the range *M* ∈ [7^2^, 13^2^], the correlation is relatively high, which allows us to use 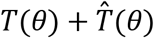 to estimate *n_c_* and *E*. Due to the complex geometrical features of hexagonal grids with different orientations, it is very difficult to completely explain the exact shape of the correlation curve. Here, our conclusions only rely on the overall high correlation value as a starting point of our theoretical analyses. Because of the high correlation, when the maximum entropy *E* is reached, the values of 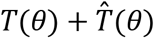 should be maximized as well at a specific value of *θ*, i.e., 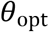. Hence, we rotate the “ + “-shape in Fig 4A to find the maximum value of 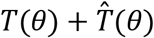 occuring at the optimal orientation *θ*_opt_. (Fig 4A shows an example *M* = 5^2^ for illustration).

**Fig 4.**
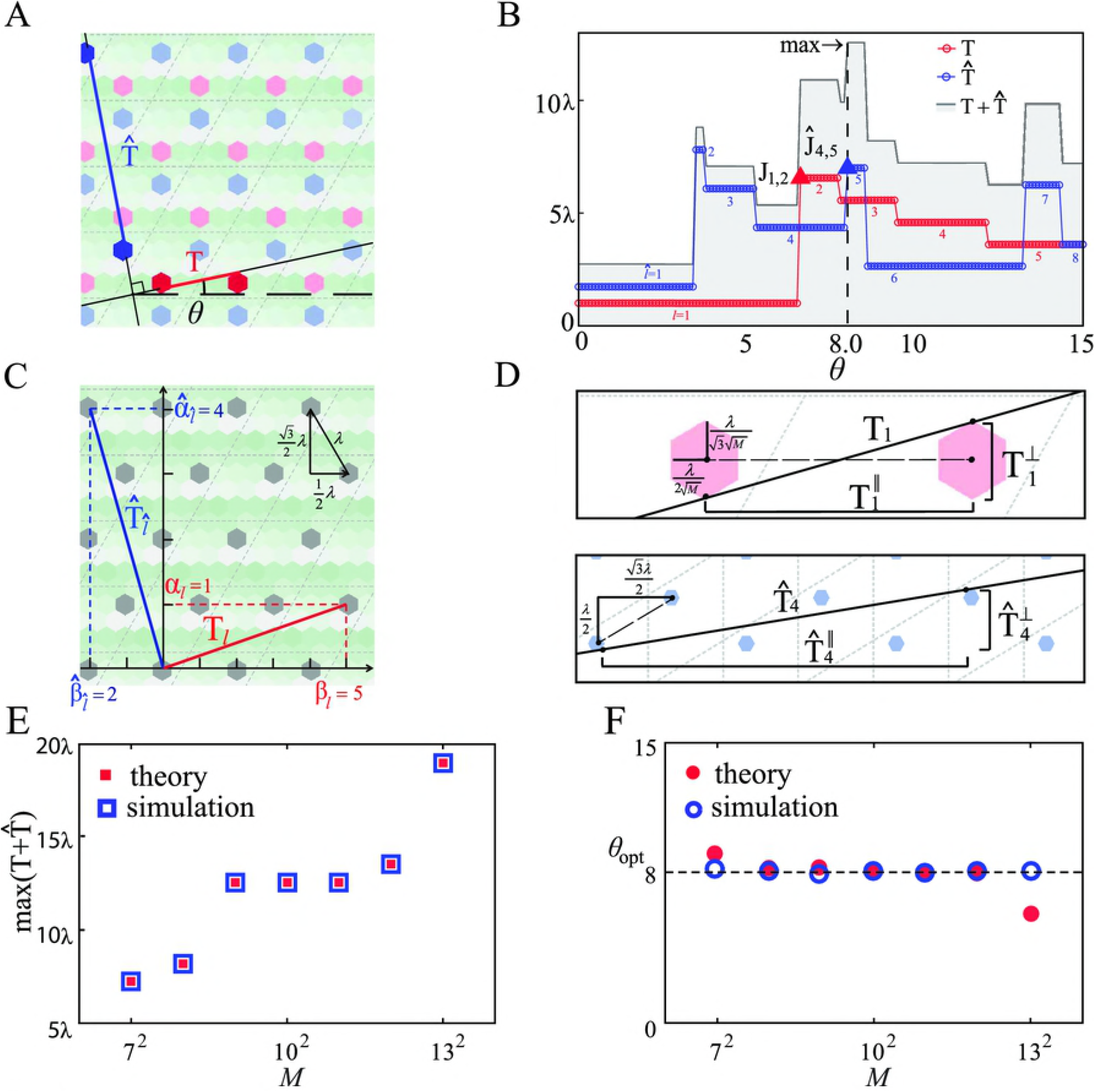
Theoretical analysis of the optimal orientation in an ideal situation. (**A**) Schematic illustration of the minimum repeating distance *T* and 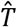 of two adjacent sites encoded by the same cell along the two orthogonal lines at orientation *θ*. (**B**) Numerical results of *T*, 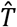 and 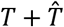 as a function of the orientation *θ* for *M* = 10^2^ showing a number of abrupt transition points and plateaus. The plateaus associated with *T* and 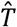 are denoted by *l* and 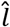, respectively, where *l* = 1,···,5 and 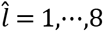, with the respective transition points as *J*_*l,l*+1_ and 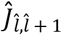. The maximum value of 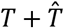 occurs at 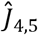. (**C**) Definitions of the unit length in the spatial encoding pattern for calculating plateau values of 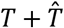. Two fundamental directions ⊤ and || are defined based on the periodic property of the spatial encoding pattern. The unit length along the fundamental directions is half of the distance between two adjacent sites encoded by the same cell. They are 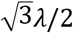 and *λ*/2 for ⊤ and || direction, respectively. *α_l_* and || denote the absolute number of unit length of the minimum length *T* in plateau *l* along ⊤ and *β*, respectively. 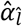 and 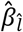 denote the absolute number of unit length of the minimum length of 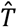 in plateau 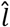 along ⊤ and ||, respectively. (**D**) Geometrical configurations of *T*_1_ and 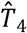, where the tangent line between two sites is associated with transition points *J*_1,2_ and 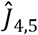. If the orientation *θ* is larger than the angle of the tangent line, *T*_1_ and 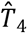 will suddenly change to *T*_2_ and 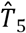, respectively. The critical orientation *θ* associated with the transition points can be obtained through 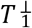 and 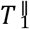, and 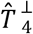 and 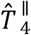, respectively. (**E**) 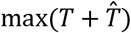 as a function of *M* obtained by simulations and theoretical analysis from Eq. (12). (**F**) Optimal orientation *θ*_opt_ as a function of *M* obtained by simulations and theoretical analysis from Eq. (8). Numerical results are obtained by identifying the orientation associated with the highest entropy *E* defined in Eq. (6). The geometrical ratio *r* is 1.5 and the environment size is 1.5 × 1.5m^2^.

Fig 4B shows (as an example for *M* = 10^2^) the numerically calculated dependence of *T* and 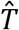 on *θ*, which exhibit a stepwise behavior with distinct plateaus and abrupt transitions between adjacent plateaus. For convenience, we define *T_l_* = *T*(*θ*) and 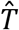, and use 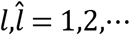 to denote the plateaus in *T* and 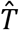, respectively, with the corresponding transition points denoted by *J*_*l,l* + 1_ and 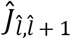, as shown in Fig 4B. (The results of *T* and 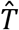 versus *θ* for *M* = 8^2^ are shown in S7A Fig as an example). It can be seen that the optimal value 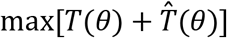 occurs at *J*_1,2_ for *M* = 7^2^,8^2^ and at *Ĵ*_4,5_ for *M* = 9^2^ to 12^2^ (S7B Fig). The transition points occur at about 8 degrees. This finding provides a base for a geometrical prediction of the value of 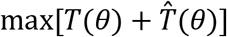 associated with different plateaus and the critical transition points among the plateaus.

Fig 4C shows the geometrical relation between *T*(*θ*) and 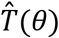 associated with plateaus and the periodic scale along the two fundamental directions (⊤ and ||) of the encoded grid. Both *T*(*θ*) and 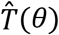 can be captured by the number 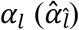 and 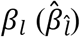 of the unit length (half of the two nearest sites encoded by the same cell) along the fundamental directions for plateau 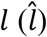. Note that the unit lengths along the fundamental directions are available, because they are only determined by *λ* in module *k*. Thus, analytical results can be obtained based on the specific geometrical feature of the plateau associated with 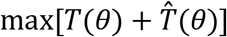.

Fig 4D show typical geometrical features at two transition points. We see that an abrupt transition occurs in the scenario when a line becomes the tangent of two nearest hexagonal sites *u* encoded by the same cell. As the line (“ + “-shape) continues to rotate in the counter-clockwise direction, *T_l_* suddenly changes to *T*_*l* +1_, indicating a discontinuous transition. The values of *T_l_* and *J*_*l,l* +1_ can be analytically calculated in terms of the geometrical features of the tangent line, as shown in Fig 4E. For deriving 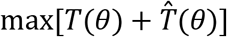, the position of tangent points can be neglected for simplicity and the distance between the centers of two neighbouring encoded sites is able to ensure prediction accuracy (see Eq. (12) and Methods). Fig 4E shows a good agreement between the analytical and simulation results of 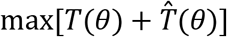. For the critical transition point associated with *θ*_opt_, the tangent points should be considered to enhance accuracy, as shown in Fig 4D. Finally, *θ*_opt_ in the region *M* ∈ [7^2^,13^2^] can be formulated in terms of the angle of the tangent:

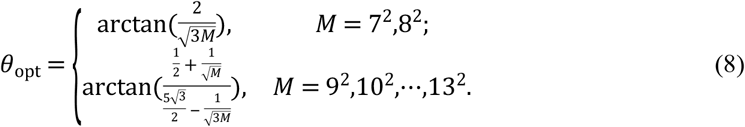

The analytical predictions of *θ*_opt_ in the absence of noise agree with simulation results in the region of *M* ∈ [7^2^,12^2^], as shown in Fig 4F (see Methods for detailed calculations).

It is noteworthy that the analytical results of *θ*_opt_ in Eq. (8) are exclusively determined by *M*, regardless of the geometrical ratio *r* of modules, which is consistent with the simulation results in Fig 3B, further validating our analysis. However, in the presence of noise, the theoretical analysis based on *T*(*θ*) is not valid anymore, because of the significant driving effect of noise on *θ*_opt_ = 8 degrees. Thus, we have to provide a new geometrical approach to dealing with noise.

### Geometrical analysis of optimal orientation in the presence of noise

We introduce a heuristic geometrical counterpart of noise to convert the effect of noise into a geometrical variable. In contrast to Fig 4A with an infinite space, here we consider a two-dimensional closed domain that can be mapped into a torus, as shown in Fig 5A [8, 35] (the torus can be obtained by connecting the upper and lower boundaries, and connecting the left and right boundaries of a square or retangular domain). The two orthogonal lines in Fig 4A at orientation *θ* become two trajectories on the surface of the torus, as illustrated in Fig 5A for three examples of *θ*. When *θ* = 0 degrees, two periodic orbits appear on the torus, whereas when *θ* = 8 or 15 degrees, the trajectories become more intricate and are not closed. The behaviors can be depicted more explicitly in mapping onto the periodic spatial unit, i.e. the rhombus in Fig 5A (see also Fig 2A). In this depiction, the two trajectories convert to two sets of lines starting from the left-bottom corner and folding onto the boundaries of the rhombus. Here ‘folding’ refers to terminating at a point at one boundary and recurring at the same point at the opposite boundary. For *θ* = 0 degrees, one line folds onto the top right corner and the other line fold onto the bottom right corner, and both return to the starting point, leading to periodic orbits. In contrast, at *θ* = 8 or 15 degrees, the folding process results in two groups of parallel lines. The key to our theory lies in quantifying the effect of noise on the basis of the folding lines in the rhombus. Fig 5B exhibits the geometrical counterpart of the noise effect exerted on a narrow belt (line). It explicitly demonstrates that the effect of noise is akin to expand the width of the belt, such that more cells are engaged in encoding and *n_c_* increases. Statistically, a stronger effect of noise is associated with a wider belt, and we use the noise-induced belt width Δ around a line to estimate the effect of noise.

**Fig 5.**
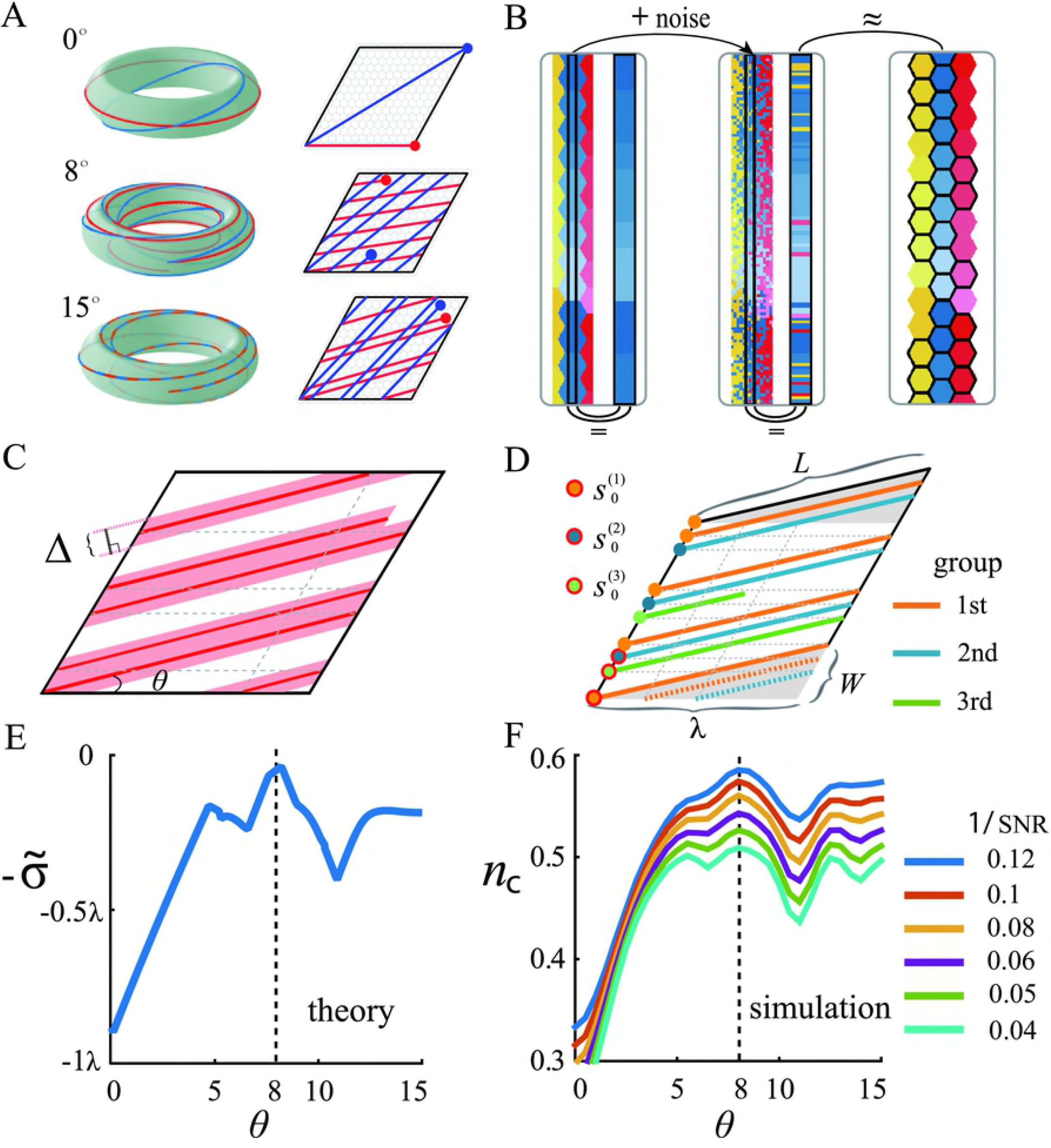
Theoretical analysis of the optimal orientation in the presence of noise. (**A**) Three toruses of a closed space and three spatial units (rhombus) on the torus for *θ* = 0, 8 and 15 degrees, respectively. The two trajectories (red and blue) on a torus and the two groups of folding lines (red and blue) in a rhombus correspond to the two orthogonal lines with finite length in a 2-dimensional space (see Fig. 4A). In a rhombus, each line starts from the bottom left corner and the ending points are highlighted. (**B**) Illustration of noise effect. Left panel: arbitrary belt and its widened image on the right hand side in the presence of noise (both are highlighted by a black frame). Middle panel: a belt and its widened image (highlighted by a black frame) with the same parameter as that in the left panel but in the presence of noise. Right panel: a wider belt without noise is associated with similar *n_c_* to that of a narrower belt with noise in the middle panel. Thus, the effect of noise is similar to the effect of widening a belt in the ideal situation without noise. (**C**) Noise-induced folding belts rooted in a set of folding lines at *θ*. The width of belts is denoted by Δ. The total length of the folding belts is *N. n_c_* is approximately determined by the area covered by the belts. (**D**) Illustration of a transformed rhombus and definitions of variables. Three groups of parallel light beams starting from the bottom left corner are displayed. The starting point of the first light beam of each group is highlighted. (**E**) Theoretical predictions of 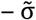 as a function of *θ* from Eq. (24). 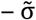 peaks at *θ* = 8 degrees, indicating *θ*_opt_ = 8 degrees associated with the highest *E* and *n_c_*. (**F**) Simulation results of *n_c_* as a function of *θ* for different effect of noise 1/SNR. In (**F**), *M* = 20^2^ and the environment size is 1.5 × 1.5m^2^.

As shown in Fig 5C, because of the effect of noise, a set of parallel lines is transformed into a set of belts with overlapping regions. As a result, *n_c_* is approximately determined by the area covered by the belts, and the larger the area, the higher the value of *n_c_*. Simulation results of the covered area as a function of *Δ* show that in a wide range of Δ, the largest area arises at *θ* = 8 degrees, substantiating our heuristic hypothesis of the geometrical counterpart of the effect of noise (see S8 Fig). Actually, the parameter *Δ* is not necessary to assess the effect of noise. The key to eliminating *Δ* lies in the fact that the area covered by the parallel belts is inversely correlated with the size of the overlapping areas. Moreover, the total overlapping area can be captured by the variance of the interval between the central lines of two nearest belts. Intuitively, a smaller variance associated with more similar intervals leads to smaller overlapping areas and a larger coverage. Thus, we analyze the standard deviation 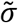 of the intervals for different *θ* to estimate *n_c_*. As shown in Fig 5D, we implement a geometrical transformation on the rhombus unit, such that all folding lines are parallel to the boundaries of the new rhombus to facilitate our analysis. This allows us to formulate the starting point of each line during the folding process by virtue of a set of inequalities (see Methods for details). The starting points are the key to deriving the theoretical result of 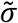, as shown in Fig 5E. Fig 5F shows simulation results of *n_c_* versus *θ* for different influences of noise, which are in good agreement with analytical predictions in Fig 5E, especially for the global maxima at *θ* = 8 degrees, the local maxima at *θ* = 15 degrees, and the local minima at *θ* = 11 degrees and 7 degrees. The analytical and simulation results provide strong evidence that an optimal orientation at 8 degrees is associated with maximum information entropy in the presence of noise.

### Reinforcement learning model for the evolution of orientation

The results described so far demonstrate the benefit of an orientation at 8 degrees for maximizing spatial information content in the presence of noise both numerically and analytically. However, how hexagonal grids gradually evolve towards this optimal orientation remains an outstanding question. We address this issue by a reinforcement learning model that is trained using experimentally recorded trajectories of rats (data:http://www.ntnu.edu/kavli/research/grid-cell-data). Broadly, reinforcement learning is a type of machine learning, which uses experience gained through interacting with the environment to evaluate feedback and improve decision making. We hypothesize that rats implement a spatial encoding scheme during exploration of a new environment and optimize their spatial representation for a higher chance of survival. More specifically, we assume that a rat is able to evaluate its orientation *θ* based on past experience and modify *θ* to maximize spatial information content.

The essential ingredients of our reinforcement learning model are an adopted strategy set, in which the evolutionary fitness of each strategy is evaluated based on experience. To be concrete, we assume that the strategy set consists of all possible orientation angles *θ_1_, *θ*_2_*, ···, *θ_n_* in the range [0,15] degrees with a small interval between two adjacent strategies. As shown in Fig 6A, we assign the same initial fitness *F*(*θ_i_*,0) to all strategies. We assume that a rat explores the environment using the same strategy in a round for a certain amount of time. In an arbitrary round, say 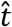, the animal chooses a current strategy 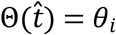 from the strategy set according to the selection probability determined by the fitness of strategy *θ_i_* at time 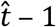 (see Eq. (26) in Methods for the definition of selection probability). After 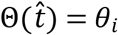 in round 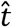 is decided, the rat’s grid cells create hexagonal grids, encode spatial locations and evaluate spatial representation with respect to its moving trajectory in round 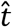. In particular, it is reasonable to assume that only locations pertaining to trajectories are encoded and used to calculate information entropy 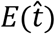 in the current round 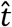 (see Eq. (28) in Methods for the definition of 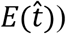. Subsequently, the fitness 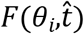 of strategy *θ_i_* used in round 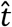 is updated according to Eq. (27), in which 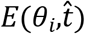 associated with *θ_i_* in all rounds prior to 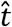 are used. In the next round 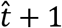, the rat repeats this process. After a number of rounds, we obtain a distribution of the frequency of strategies (see Methods for more details).

**Fig 6.**
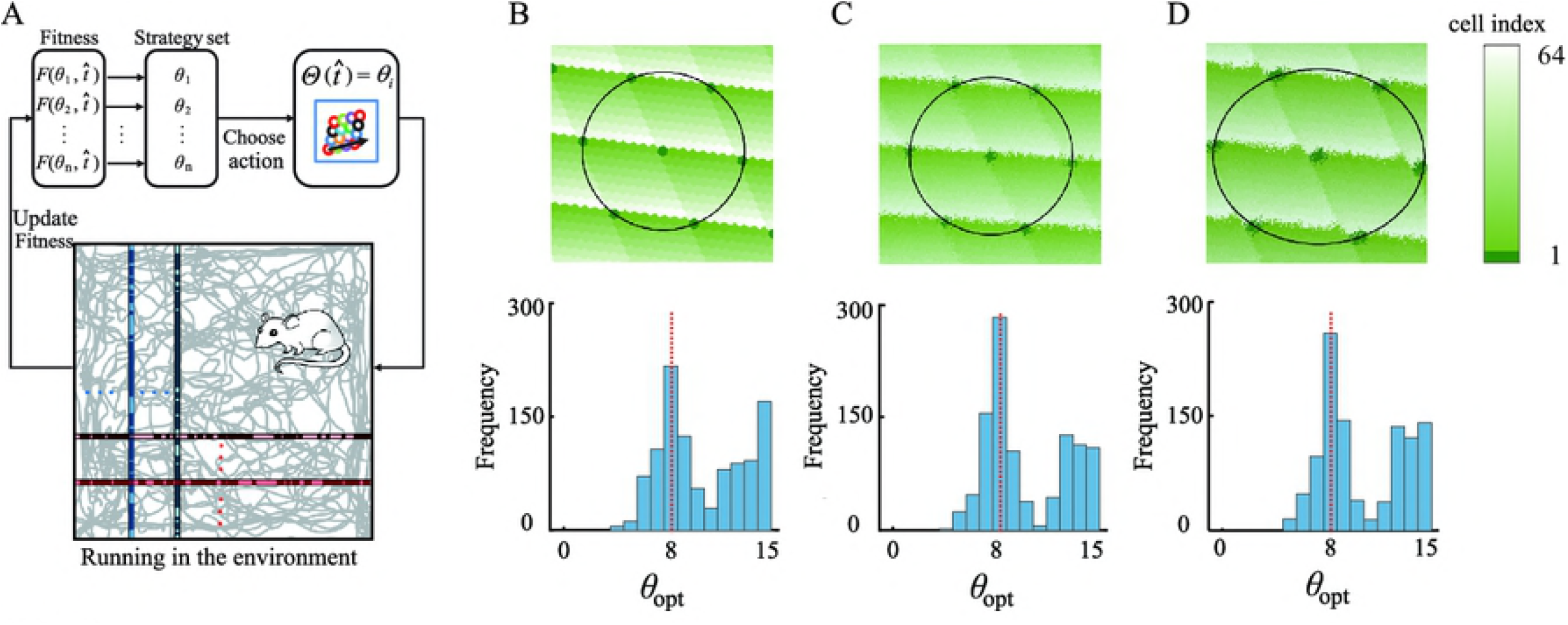
Reinforcement learning model for the evolution towards the optimal orientation. (**A**) Schematic illustration of the reinforcement learning model, where the current strategy Θ used by rats is determined by Eq. (26) and the strategy fitness *F* is updated according to Eq. (27). The trajectories of rats in experiments are used to compute spatial entropy over all passing locations in all belts in both directions. (**B-D**) Frequency of using strategy *θ* in the ideal situation without noise and elliptic distortion (**B**), in a situation with noise but without distortion (**C**) and in a realistic situation with both noise and elliptic distortion (**D**). The parameters in the reinforcement learning models are *r* = 1.44, *M* = 10^2^, *t* = 500, *w*(*t*) = *e^κt^* and *κ* = 0.15, environment size 1.5 × 1.5m^2^ and *ε* = 1.17. The frequency distributions are obtained by 1000 independent realizations. In (**C**) and (**D**), the effect of noise 1/SNR = 0.08. In all simulation, *f*_max_ = 200Hz.

Fig 6B-D show, respectively, three sets of representative results from the reinforcement learning model: in the ideal situation, in the presence of noise only and in the presence of both noise and elliptic distortion. A common phenomenon is that there are two peaks in the frequency distributions at *θ* = 8 degrees and about 15 degrees, respectively, where the former is the global maxima and the latter is a local maxima. The global maxima at *θ* = 8 degrees is consistent with recent experimental observations. Meanwhile, the local maxima at *θ* = 15 degrees is also supported by some previous experimental evidence [15]. We speculate that the experimental observation of *θ* = 15 degrees may be attributed to a relatively small amount of recorded grid cells in the experiments, leading to a bias towards the local maxima at *θ* = 15 degrees. The distribution obtained by our reinforcement learning model provides a possible explanation for the different experimental results in the literature.

Due to finite evolutionary time assumed in our reinforcement learning model, the strategy pool still consists of a rich variety of strategies in spite of the advantage of *θ* = 8 degrees. In principle, a strategy with evolutionary advantages possesses higher fitness, and the “strong gets stronger” effect during the process reinforces the strategy. If the evolutionary time is sufficiently long, the optimal strategy *θ* = 8 degrees will prevail eventually and dominate the strategy set, and all the other strategies will be eliminated. To be consistent with experiments, our results pertain to temporal stages (see Methods for more details) rather than the thermodynamic limit. As a result, a distribution of strategies with both the global and a local maxima are obtained.

## Discussion

Converging experimental evidence has revealed that the main axes of grid cells are anchored to environmental borders at an offset of about 8 degrees. Here, we aimed at understanding the possible functional roles (i.e., the evolutionary advantage) and the physiological mechanisms underlying this phenomenon.

By combining a computer simulation of the spatial coding properties of the grid cell network with analytical considerations and a reinforcement learning model, we found evidence that a grid axis offset of 8 degrees maximizes the spatial information content specifically in thre presence of noise.

We first implemented a computer simulation of the grid cell network and analyzed the influence of grid orientation on the grid cell network’s spatial coding properties, depending on various factors such as grid cell numbers, the effect of noise, the spatial ratio of module scales, and elliptic distortions. Unexpectedly, we found that a grid orientation of 8 degrees maximizes spatial information content across a wide range of conditions specifically in the presence of noise. In a noise-free situation, an orientation at 8 degrees is only optimal within a narrow and putatively unrealistic range of the numbers of grid cells, beause the maximum number in that range is smaller than the putative number of grid cells based on experiments [15]. By contrast, in the presence of noise, an orientation of 8 degrees maximizes spatial information as long as the number of grid cells exceeds some lower bound. This lower bound is small, insensitive to noise, and largely independent of other internal and external conditions. These results demonstrate that noise plays a significant role in driving optimal orientation towards 8 degrees and mammals optimize spatial coding in the presence of noise. Although noise has been known as a driving force of many biological phenomena [43], e.g. genetic mutation, the effect of noise on shaping the hexagonal firing pattern of grid cells has not be revealed before.

To unveil the mechanisms underlying the noise-induced universal orientation, we provide two theoretical analyses for the ideal and noisy situations. A property of information entropy is exploited to enable analytical predictions of the optimal orientation, i.e., the strong correlation between information entropy and the number of grid cells. For the ideal situation, the number of encoded grid cells along a line is estimated by the minimum repeating distance between two nearest sites encoded by the same cell. For the realistic situation with noise and elliptic distortion, the effect of noise is akin to transforming a line to a belt with a certain width. The analytical predictions are in good agreement with simulation results across a wide range of grid cell numbers and noise effects.

Finally, we modeled the temporal evolution of grid cell orientations towards the optimal angle via a reinforcement learning model based on experimentally observed behavioral trajectories of rats. During the time-course of reinforcement learning, the model evaluates the fitness (i.e., information entropy along trajectories) depending on various strategies that are defined by sets of orientation angles. If an orientation angle leads to a higher information entropy prior to a moment, the angle will be adopted with a higher probability at that moment. As the process continues, all possible strategies (angles) will be employed, but the strategies with higher fitness will be used more frequently. Our results demonstrate that for both ideal and realistic situations, the likelihood of strategies exhibits a global maximum at 8 degrees and a second maximum at about 15 degrees, in line with experimental evidence [15, 23-24].

The main hypothesis of our coding scheme – i.e., that the orientation of grid patterns maximizes spatial information – has been partially validated by a very recent experiment in humans [44]. In the experiment, two axes with different amounts of information are arranged in a circular environment. It is found that grid cell systems reduce the uncertainty of representation by aligning grid patterns to the axis of greatest information. The concept of uncertainty reduction is analogous to the maximization of spatial information in our coding scheme, despite the very different experimental settings. The orientation aligned to the axis of greatest information, suggesting a similar effect as for the optimal orientation at 8 degrees in a square environment. Taken together, these recent experimental results substantiate the idea that grid orientation serves to maximize spatial information.

It is worth noting that our encoding model can not only explain the functional benefit of the empirically observed grid orientation but also generates experimentally testable predictions on the lower bound of cell numbers *M* in each module. According to the results in Fig 3K, this lower bound is determined by the effect of noise measured by the inverse of the signal to noise ratio 1/SNR. The effect of noise that depends on the duration time *τ* can be estimated from hexagonal patterns in any one module documented in experiments (see S2 Methods). As shown in S9 Fig, *τ* estimated in experiments is about 0.19s and the effect of noise can be assessed to be 0.16. According to Fig 3K, the minimum cell numbers *M* for inducing an optimal orientation at 8 degrees can be predicted to be 16^2^ in each module. It is noteworthy that the actual number of grid cells may be larger than this prediction. In fact, the result predicts that under a certain amount of noise, the number of grid cells should be larger than a critical value to ensure an optimal encoding at an orientation of 8 degrees. If the amount of noise is increased by reducing *τ*, in order to achieve an valid coding, more grid cells are needed. This is explicitly reflected by the increasing trend in Fig 3K as *τ* increases. Since the exact number of grid cells currently cannot be empirically determined, the model’s predictions have implications in constraining future modeling approaches.

Besides square environments, our coding model can be easily applied to other environments with regular shapes. For example, in a rectangular environment, we vary the ratio of the longer edge to the lower edge, and compute optimal orientation angles under different ratios. Due to the destruction of symmetry in a rectangle compared to a square, the range of *θ* becomes [0, 30] degrees. Our results show that spatial information is maximized at either *θ* = 8 degrees or *θ* = 22 degrees, depending on both the effect of noise 1/SNR and the cell numbers *M* of grid cells in each module (see S10A-B Fig). The range of the lower bound of the cell numbers *M* is quite similar to that in square environments. Note that 8 degrees and 22 degrees are symmetric values around the center of 15 degrees within the range [0, 30] degrees. When the edge ratio equals one (the rectangle becomes a square), 8 degrees and 22 degrees are completely equivalent because of the recovery of the broken geometrical symmetry. Therefore, there is an essential relationship between the orientation in a square and in a rectangular environment according to our prediction, resulting from the geometrical similarity between the two shapes. The analytical results based on the folding in a rhombus are applicable for rectangular environments as well, and the analytical results are in good agreement with simulation results based on information entropy (see S10C Fig). The numerical and theoretical predictions in rectangular environments could be validated by future experiments.

It has been argued that grid cells play a crucial role for path integration, a navigation strategy that is based on an idiothetic reference frame [8-10, 45-46]. As a consequence, an orientation of grid patterns that maximizes spatial information may benefit path integration. Specifically, during path integration, several types of information are integrated, such as spatial locations, speed, and directions. In addition to the putative role of place cells, speed cells and head directional cells for path integration, grid cells can provide necessary information as well. On the one hand, grid cells in MEC may input lowdimensional information into different high-dimensional cells, e.g., place, speed and head directional cells, to implement path integration. On the other hand, during path integration, errors from noise will be aggregated in the absence of salient landmarks or boundaries, and place fields will drift [29]. The grid cell systems may help reduce the errors because the low-dimensional cognitive map created by grid cells is relatively stable across different contexts and environments. Note that to precisely encode either distance [35] or speed during path integration, an accurate self-localization is imperative. The orientation of hexagonal grid patterns maximizes spatial information and discriminates different locations in an optimal manner, so that the tracking of distance and speed is favored by this orientation as well may improve path integration.

Many open issues remain for a more general understanding of the dynamics and functions of grid cells during spatial navigation. For example, how does the activity of place cells in the hippocampus relate to the orientation of grid cells [47-50]? How do grid cells modulate the coding scheme for two separate spaces when the two spaces are being merged into a single space [51-52]? Our modeling approach consisting of a spatial encoding scheme, a method to characterize encoding capacity, geometrical analyses for both ideal and realistic situations, and a reinforcement learning model offers tools for addressing these issues to unravel the mysteries of the navigation ability of the mammalian brain.

## Methods

### Spatial firing rate of grid cells

The idealized firing rate 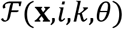 of cell *i* in module *k* with orientation *θ* at an arbitrary location (**x**) is

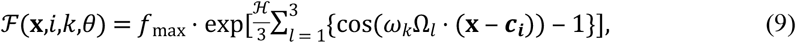

where **x** = (*x,y*) ∈ ℝ^2^ denotes the location, Ω_*l*_ = (cos(*φ_l_*),sin(*φ_l_*)) with *φ_l_* =-*π*/6 + *lπ*/3 + *θ* for *l* = 1,2,3, *ω_k_* = 2*π*/[*λ*(*k*)sin(*π*/3)], 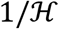 measures the cell’s relative tuning width, *f*_max_ is the maximum expected firing rate, ***c**_i_* is the spatial phase of cell *i* (any one of the centers of the firing fields - see Fig 1). Because of the property of modularity, we assign the same tuning function to each grid cell within a module so that they have the same *f*_max_ and 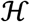 values (e.g., *f*_max_ = 200 and 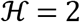), but the values of the spatial phase parameter ***c*** are equidistantly arranged, as shown in Fig 2A.

### Spatial noise in firing intensity

With respect to measurement noise and intrinsic noise in firing rate, the actual firing intensity *I*(**x**,*i,k,θ*) would scatter around the ideal (expected) value 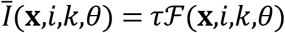. *I*(**x**,*i,k,θ*) at location *x* is assumed to follow a Poisson distribution with expected value 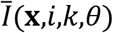 as used in (*14,18–20*):

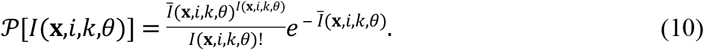

### Elliptic distortion

Suppose that the firing field centers of a grid cell lie on an ellipse with a semi-major axis *a*_1_ and semiminor axis *a*_2_ with *a*_1_ > *a*_2_ (see the yellow axis *a*_1_ and black axis α_2_ in the ellipse in Fig 3K). To model ellipticity, the wave vectors in Eq. (9) are rescaled to Ω_*l*_ = (cos(*φ_l_*)/*ε*,sin(*φ_l_*)), where *ε* denotes the elliptic intensity and is defined as the ratio of the major axis of the ellipse to the minor axis: *ε* = *a*_1_/*a*_2_.

### Geometrical analysis in the ideal situation

The value of 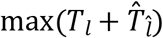 can be estimated by the length of the tangent at *θ*_o*pt*_ associated with a specific plateau (see Fig 4E). Note that the geometry of tangent points has negligible effect on 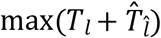 because of the plateaus. We thus only calculate the interval between the centers of two nearest hexagonal sites encoded by the same cell pertaining to the tangent (Fig 4E). In general, *T_l_* and 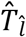 of plateau *l* and 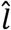 can be captured by virtue of the two fundamental directions (⊤ and ||) of the grid (see Fig 4D):

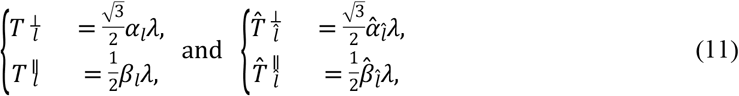

where *α_l_*, *β_l_*, 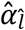, and 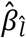 are the numbers of the unit length (half distance between two nearest sites encoded by the same cell) along the two fundamental directions for plateaus *l* and 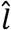, respectively (see Fig 4D). Corresponding to the maximum value of *E*, the value of 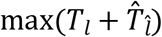 can be estimated as

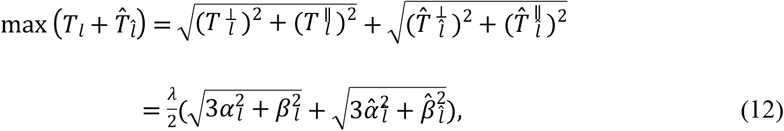

where 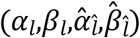 are (1,9,3,1) for *M* = 7^2^, (1,11,3,1) for *M* = 8^2^, (1,11,8,2) for *M* = 9^2^ to 11^2^, (1,13,8,2) for *M* = 12^2^, and (1,17,12,2) for *M* = 13^2^. The analytical results are in good agreement with simulation results, as shown in Fig 4F.

To obtain the value of *θ*_opt_ determined by the tangent angle, the geometrical features of the tangent points can increase the prediction accuracy (see Fig 4E). We formulate all the transition points. There are two kinds of transitions: from a lower plateau to an upper one and vice versa. For the former class, i.e., *T_l_* < *T*_*l* + 1_ (or 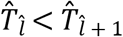), the geometrical relation in Fig 4E gives

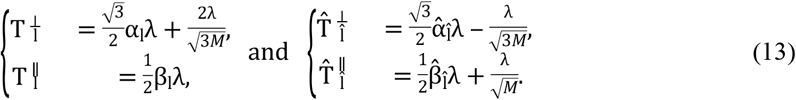

For the second type, i.e., *T_l_* < *T*_*l* + 1_ (or 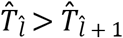), the tangent behavior is different from that in Fig 4E, where the tangent point is at the top of the left field and at the bottom of the right field. In this case, we have

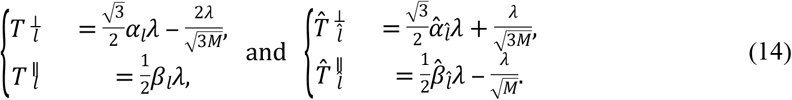

The quantities *θ*_*l,l* + 1_ and 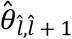 can be obtained as 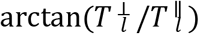 and 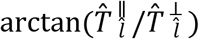, respectively.

The value of *θ*_opt_ can be obtained from the tangent of two nearest sites encoded by the same cell. In particular, for *M* = 7^2^,8^2^, the value of *θ*_opt_ is determined by *J*_1,2_. Using the values of *α*_1_ and *β*_1_ for *l* = 1 (*α*_1_ = 0,*β*_1_ = 2), we have *θ*_opt_ = *θ*_1,2_. For *M* = 9^2^ to 13^2^, the value of *θ*_opt_ is determined by 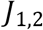. Using the values of 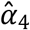 and 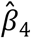 for 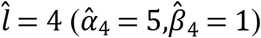, we have 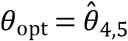. Finally, we obtain Eq. (8) for *θ*_opt_ in the region *M* ∈ [7^2^,13^2^].

### Geometrical analysis in the presence of noise

We present the process of theoretically deriving the starting points of folding lines on the left boundary of the transformed rhombus in Fig 5D.

We define the bottom left corner to be the zero point, and suppose that there is a beam of light emitted from the zero point along direction *θ* in the rhombus unit, as shown in Fig 5C. If the light reaches the right (top) boundary, it will return to the same point in the left (bottom) boundary because of the closed boundary condition. The representation is able to capture the scenario that a line goes through a number of rhombus units on the surface of the encoded torus in Fig 5A. The other trajectory on the surface of the torus (orthogonal line in Fig 4A) is a beam of light along 30 – *θ* degrees in the rhombus unit. Below we focus on the analysis associated with *θ*.

To simplify our analysis, we transform the rhombus in Fig 5C to that in Fig 5D by cutting the bottom region (grey region below the light beam starting from the zero point) and moving it to the top. In the new rhombus, all light beams are parallel to the top and bottom boundaries. The first beam emitted from the zero point (starting point 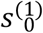) overlaps with the bottom boundary. After it reaches the right-bottom corner point, it will return to a new starting point on the left boundary with a fixed distance *W* from the zero point. This process is repeated until a starting point is reached that is higher than the top boundary. The set of light beams, the starting points of which are lower than *λ* is defined to be the 1st group. The first light beam that exceeds the top boundary returns to bottom with a starting point at 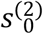 on the left boundary according to the periodic boundary condition, and it becomes the first beam of the 2nd group of beams, as shown in Fig 5D. We iterate this process until the total length *N* of the beams are reached, and several groups arise. Within each group, the interval between two nearest beams is always *W*, such that if the starting point of the first beam of each group is identified, all starting points of beams can be calculated analytically in a group. Thus, our first task is to derive the starting point 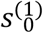 of the first beam and the number of beams in the 1st group, which can be solved via

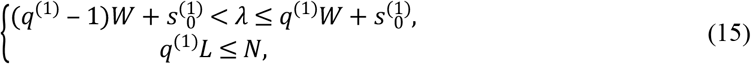

where *q*^(1)^ is the number of beams in the first group, 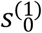 is the starting point of the first beam in the first group (zero point here), *L* = *λ*sin(120°)/sin(60° – *θ*) is the length of the top (bottom) boundary in the transformed rhombus and *W* = *λ*sin(*θ*)/sin(60° – *θ*) is the interval between two beams long the left (or right) boundary. The first inequality is formulated based on the boundary condition to characterize that the *q*^(1)^th beam is below the top boundary but the (*q*^(1)^ + 1)th beam exceeds the top boundary. The second inequality describes that the total length of beams in the 1st group should not exceed *N*. The two inequalities can exclusively yield the number *q*^(1)^ of beams in the 1st group:

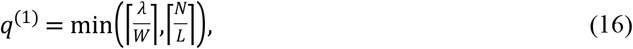

where ⌈ · ⌉ denotes rounding up to an integer. Thus, the starting point s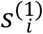 of an arbitrary beam, say *i* in the 1st group can be written by

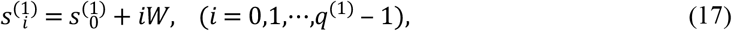

where the subscript *i* denotes the *i*th beam and 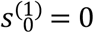. Akin to the 1st group, the starting points of beams in all groups can be solved iteratively via a general form of inequalities

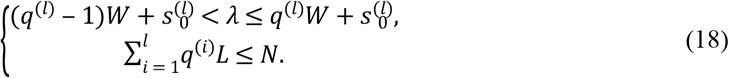

The number *q*^(*l*)^ of beams in the *l*th group is:

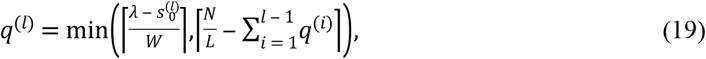

where

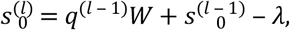

and the starting points of the *l*th group are

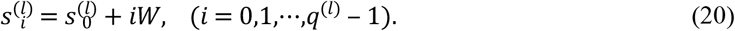

We thus obtain a set *S* composed of all the starting points in all groups:

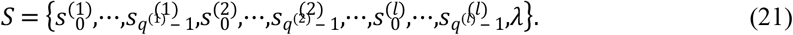

We sort the elements of *S* in ascending order, yielding

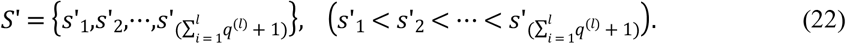

Thus, the interval *ς_i_* of two nearest beams is

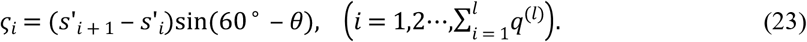

Then the standard deviation *σ*_1_ of *ς_i_* along direction *θ* can be calculated via

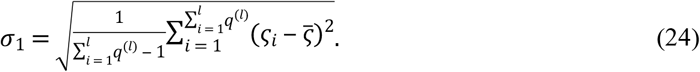

*σ*_2_ of the other groups of beams along 30° –*θ* can be analytically obtained in the same manner. The average standard deviation 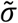 over two groups of beams is

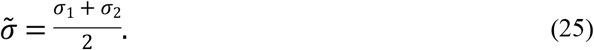

The analytical result of 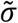 is shown in Fig 5E.

### Reinforcement learning

In an arbitrary round, say 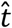, a rat chooses a current strategy 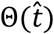 from the strategy pool according to the probability determined by its fitness values prior to 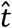 through a softmax distribution:

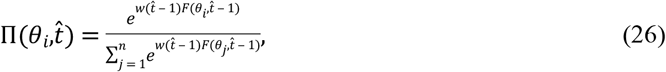

where factor *w*(*t*) = *e^κt^* with *κ* = 0.15 ensures asymptotic stability of the evolutionary process. The fitness 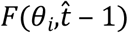 of strategy *θ_i_* in round 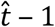 is calculated using all the rounds prior to round 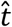:

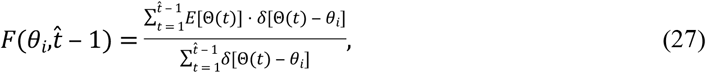

where *δ* is the Dirac function with *δ*(*l*_1_ – *l*_2_) = 1 if *l*_1_ = *l*_2_ and *δ*(*l*_1_ – *l*_2_) = 0 otherwise, *E*[Θ(*t*)] is the spatial information entropy associated with strategy Θ(*t*) in round *t*. Here the definition of *E*[Θ(*t*)] slightly differs from that in the static encoding scenario. Specifically, only locations pertaining to the trajectories of rat are used to evaluate the spatial encoding in a round. We build a database composed of 40 trajectories of rats, each of which is recorded within 10 minutes (data from http://www.ntnu.edu/kavli/research/grid-cell-data). In an arbitrary round *t*, we randomly choose a trajectory from the database, and compute *E*[Θ(*t*)] with respect to the trajectory. In the definition of *E*, *p*(*i*|0,*k,x*) is changed to

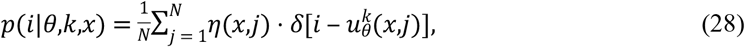

where *j* is the index of a box on the *x*th belt, *δ* is the Dirac function and *η*(*x,j*) = 1 only if location (*x,j*) pertains to the trajectory in the round (*η*(*x,j*) = 0 if (*x,j*) does not pertain to the trajectory). Analogously, *p*(*i*|*θ,k,y*) is defined in the same way but substituting 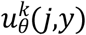 for 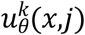, and *η*(*j,y*) for *η*(*x,j*).

## Acknowledgments

DC was supported by the Fundamental Research Funds for the Central Universities. WXW was supported by NSFC under Grant No.71631002. YCL would like to acknowledge support from the Vannevar Bush Faculty Fellowship program sponsored by the Basic Research Office of the Assistant Secretary of Defense for Research and Engineering and funded by the Office of Naval Research through Grant No. N00014-16-1-2828. NA received funding via the DFG (SFB 874/B11, SFB 1280/A02, and AX82/3).

## Author contributions

**Conceptualization:** Wen-Xu Wang, Nikolai Axmacher.

**Formal analysis:** Liang Wang, Zhanjun Zhang, Ying-Cheng Lai, Nikolai Axmacher, Wen-Xu Wang.

**Funding acquisition:** Zhanjun Zhang, Liang Wang, Wen-Xu Wang

**Investigation:** Dong Chen, Kai-Jia Sun, Liang Wang, Zhanjun Zhang, Ying-Cheng Lai, Nikolai Axmacher, Wen-Xu Wang.

**Methodology:** Dong Chen, Kai-Jia Sun, Liang Wang, Zhanjun Zhang.

**Supervision:** Nikolai Axmacher, Wen-Xu Wang.

**Visualization:** Dong Chen, Kai-Jia Sun, Liang Wang.

**Writing - original draft:** Wen-Xu Wang, Ying-Cheng Lai

**Writing - review & editing:** Nikolai Axmacher, Wen-Xu Wang

DC and KJS contribute equally, and WXW and NA contribute equally.

## Supporting information

**S1 Methods. Hamming distance.** To provide additional evidence to validate our encoding scheme, we propose a measurement based on Hamming distance for encoding capacity as an alternative to information entropy. Note that a basic requirement for any positioning task lies in the ability to distinguish vectors **u**(**x**) after the grid cell system maps the environment using orientation *θ*.

Consider two arbitrary locations **x** and **y**. They cannot be distinguished if **u**(**x**) = **u**(**y**). In addition, if the two vectors **u**(**x**), **u**(**y**) are approximately equal with a negligible difference, it would be difficult to separate them. The two locations may be conceived of as the same by the animal. The difference between **u**(**x**) and **u**(**y**) can be conveniently measured by the Hamming distance defined as:

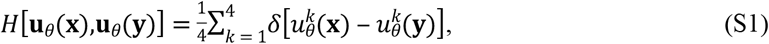

where *δ* is the Dirac function with *δ*(*l*_1_ – *l*_2_) = 1 if *l*_1_ = *l*_2_, and *δ*(*l*_1_ – *l*_2_) = 0 otherwise (see S3 Fig). If **u**(**x**) and **u**(**y**) are exactly the same, *H* is zero; if the encoded site **u**(**x**) differ from **u**(**y**), we have *H* =1. A larger value of *H* indicates a better spatial encoding and less ambiguity. For orientation *θ*, the mean Hamming distance 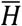 over all belts along direction **x** and **y** in a space is defined as

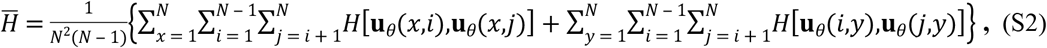

where *x*th or *y*th belt is divided into *N* unit boxes, each being denoted by index *i* or *j*. To obtain the mean Hamming distance 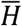, the average *H* of every belt should be obtained by averaging over *N*^2^ (*N*-1) pairs of boxes in a belt. Then 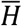 is obtained by averaging over all belts along the two directions, respectively. The results of 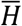 are shown in S4 Fig.

**S2 Methods. Experimental assessments of the signal to noise ratio and its influence on the predicted minimum number of grid cells *M*.** As described in the main text and shown in Fig 3, the minimum number of grid cells is crucially influenced by noise. Specifically, in case of a higher noise level, a larger number of grid cells are required for obtaining an optimal orientation at 8 degrees. Note that this effect of noise is determined by the duration *τ* across which the signal is integrated and the intrinsic firing rate. The relationship of these two factors is determined by the ratio of the variance 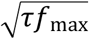 to the mean value *τf*_max_ of the Poisson distribution of firing, i.e., 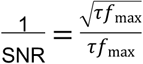, where *τf*_max_ is SNR *τf*_max_ the maximum firing intensity that measures the strength of signal. Thus, estimating the minimum value of *M* requires quantifying *τ* from experimental data, which is possible by measuring the duration of firing at a specific location. This duration depends on the time that a rat stays at a given location – i.e., since animals are moving in an environment, *τ* corresponds to the cumulative time of traversing a specific location. Now, the minimal spatial distance between pairs of spikes of grid cells defines the minimum spatial resolution of a rat. Based on these numbers, we constructed distributions of distances between any two firing locations of the grid cell, and defined the top 0.5% of smallest distances to be the scale of spatial units. As shown in S9 Fig, the average of spatial unit from all spatial patterns of grid cells is about 1.8cm, and the mean number of spatial units containing trajectories is 3120. Note that the average duration of each trajectory is 10 minutes (600s). Thus, the average duration *τ* at every spatial unit is 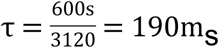 and the effect of noise 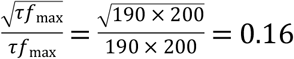 According to Fig 3K in the main text, we thus estimate the minimum number of grid cells to be *M*=16^2^ in each module.

**S1 Fig.**
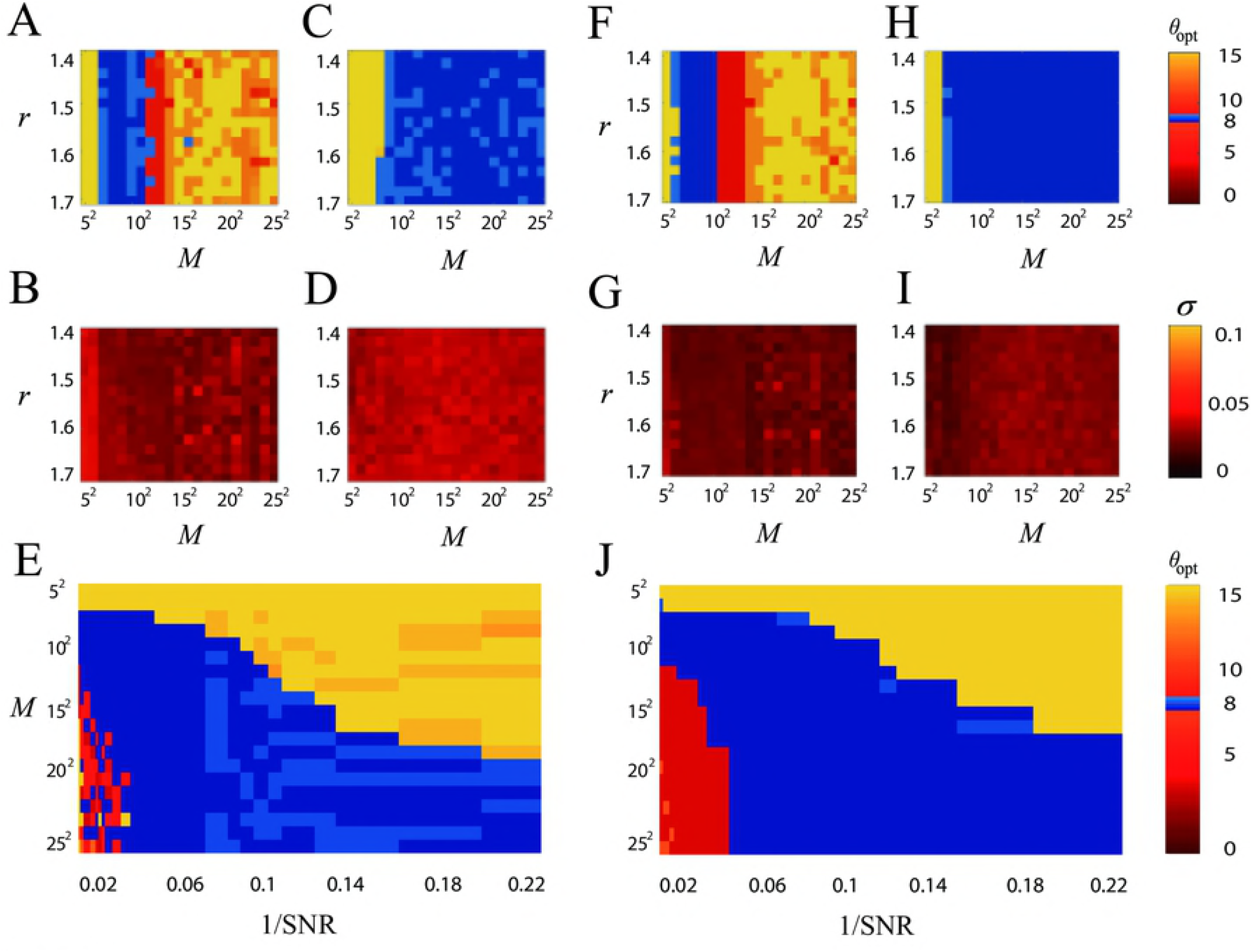
Optimal orientation associated with highest spatial information entropy for two environmental sizes. (**A**) Optimal orientation *θ*_opt_ associated with highest *E* as a function of both geometrical ratio *r* and *M* in the absence of noise. (**B**) Standard deviation *σ* of *E* of different belts at *θ*_opt_ as a function of both *r* and *M* in the absence of noise. (**C**) *θ*_opt_ as a function of both *r* and *M* in the presence of noise. (**D**) Standard deviation of *E* of different belts at *θ*_opt_ as a function of both *r* and *M* in the presence of noise. (**E**) *θ*_opt_ as a function of both noise 1/SNR and *M* in the presence of noise. In (**A**)-(**E**), the environmental size is 1.2× 1.2m^2^. The color bar of (**A**) and (**C**) represents *θ*_opt_. The color bar of (**B**) and (**D**) represents standard deviation *σ*. In (**E**) geometrical ratio *r* = 1.5. (**F**)-(**J**) are the same as (**A**)-(**E**) but the environment size is 2× 2m^2^. In (**C**) and (**H**), the effect of noise 1/SNR = 0.08. In all simulation, *f*_max_ = 200Hz.

**S2 Fig.**
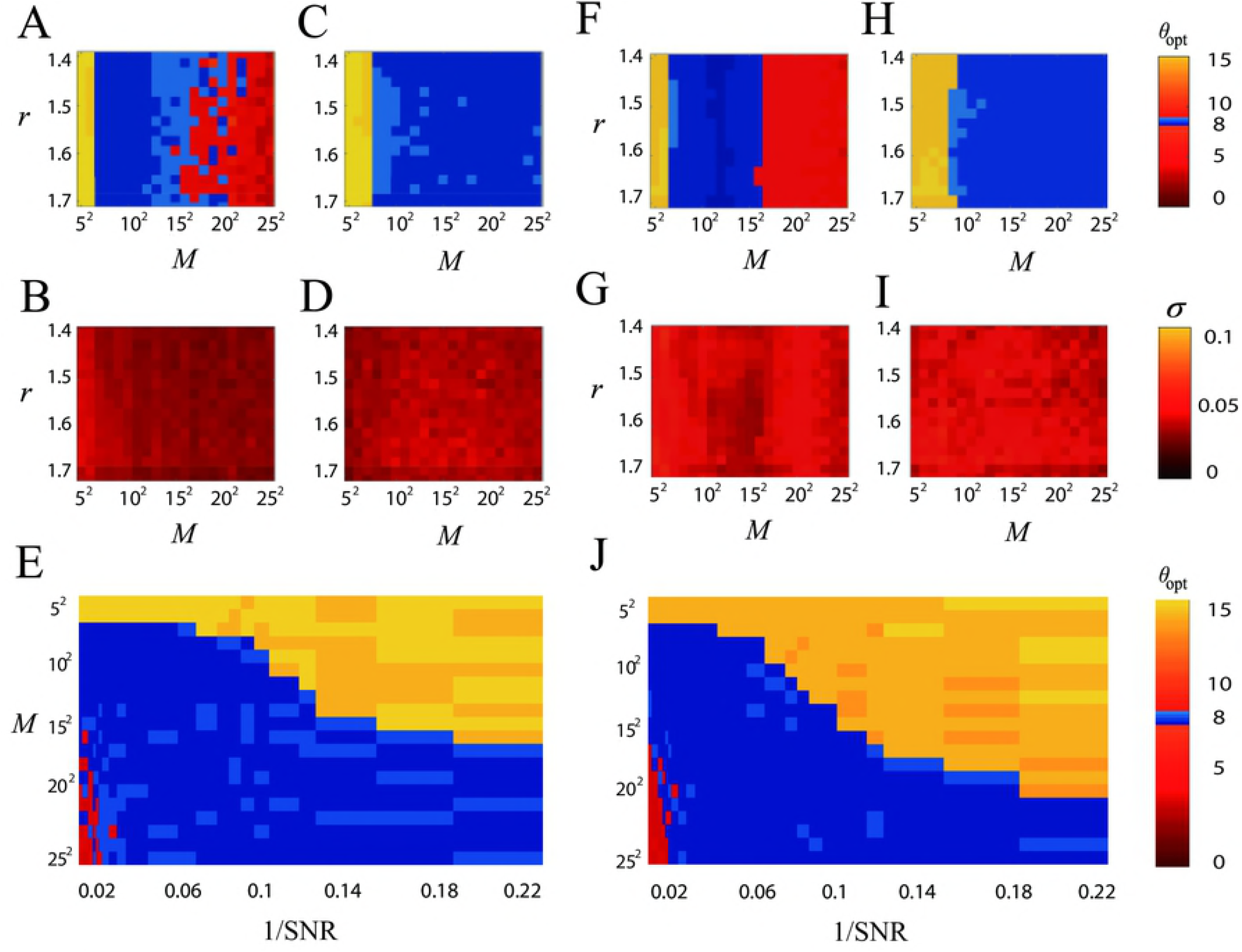
Optimal orientation associated with highest spatial information entropy for two elliptic grids. (**A**) Optimal orientation *θ*_opt_ associated with highest *E* as a function of both geometrical ratio *r* and *M* in the absence of noise. (**B**) Standard deviation *σ* of *E* of different belts at *θ*_opt_ as a function of both *r* and *M* in the absence of noise. (**C**) *θ*_opt_ as a function of both *r* and *M* in the presence of noise. (**D**) Standard deviation of *E* of different belts at *θ*_opt_ as a function of both *r* and *M* in the presence of noise. (**E**) *θ*_opt_ as a function of both noise 1/SNR and *M* in the presence of noise. In (**A**)-(**E**), the intensity of elliptic distortion is *ε* = 1.17. The color bar of (**A**) and (**C**) represents *θ*_opt_. The color bar of (**B**) and (**D**) represents standard deviation*σ*. In (**E**) geometrical ratio *r* = 1.5 and the environmental size is 1.5 × 1.5m^2^. (**F**)-(**J**) are the same as (**A-E**) but the elliptic distortion is *ε* = 1.25. In (**C**) and (**H**), the effect of noise 1/SNR = 0.08. In all simulation, *f*_max_ = 200Hz.

**S3 Fig.**
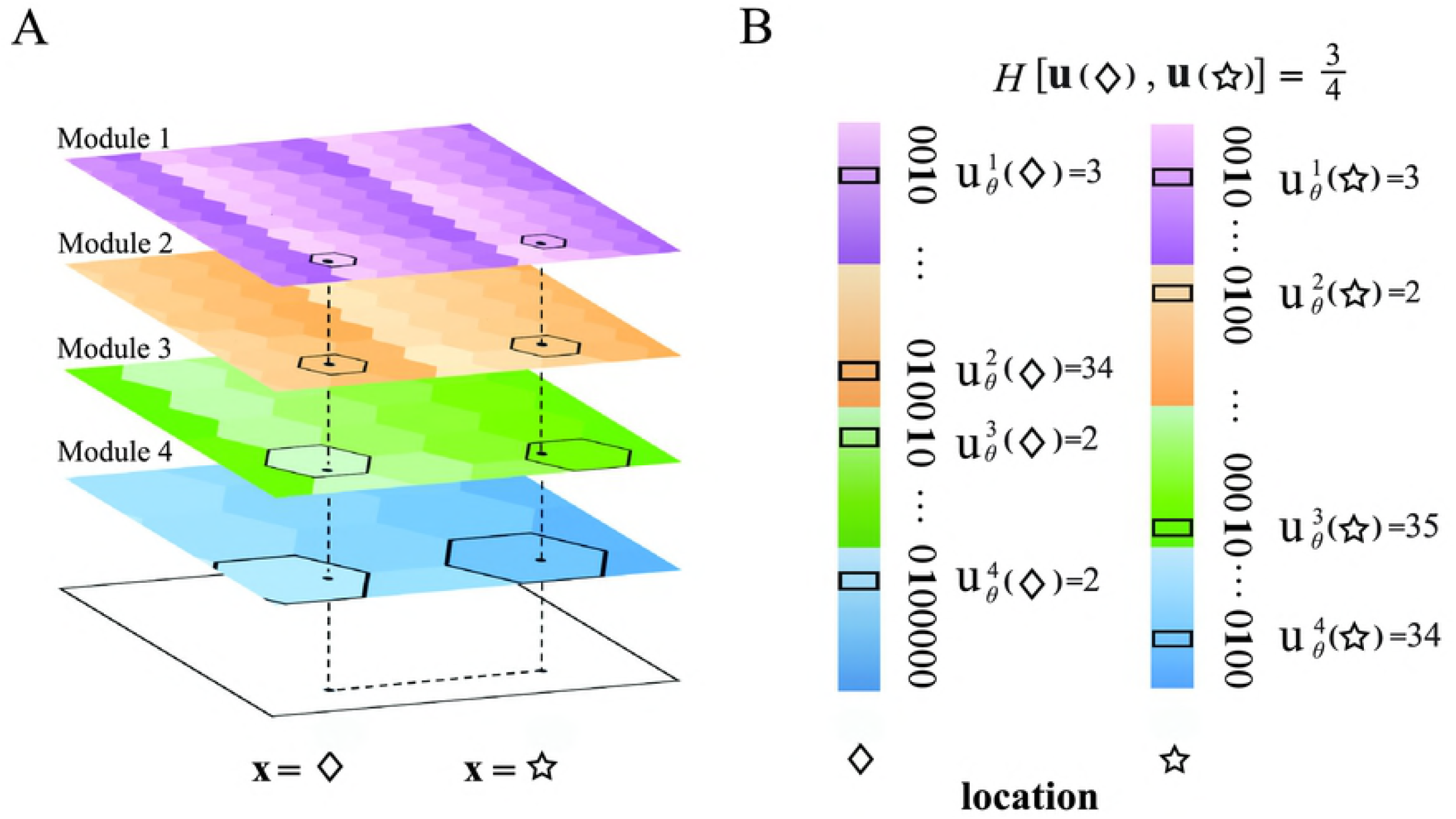
Encoding scheme of multiple modules and Hamming distance. (**A**) Hexagonally encoded sites of grid cells with different colors in modules 1 to 4. The encoded sites are distributed periodically in every module. The two locations marked by a diamond and a star are encoded by a site in each of four modules. (**B**) The spatial vector **u** of each location consists of 4 elements. If a location **x** belongs to the site encoded by cell *i* (highlighted in each module), the element in **u**(**x**) is *i*. The Hamming distance *H* of these two locations is 3/4 because there are 3 different elements between the two spatial vectors.

**S4 Fig.**
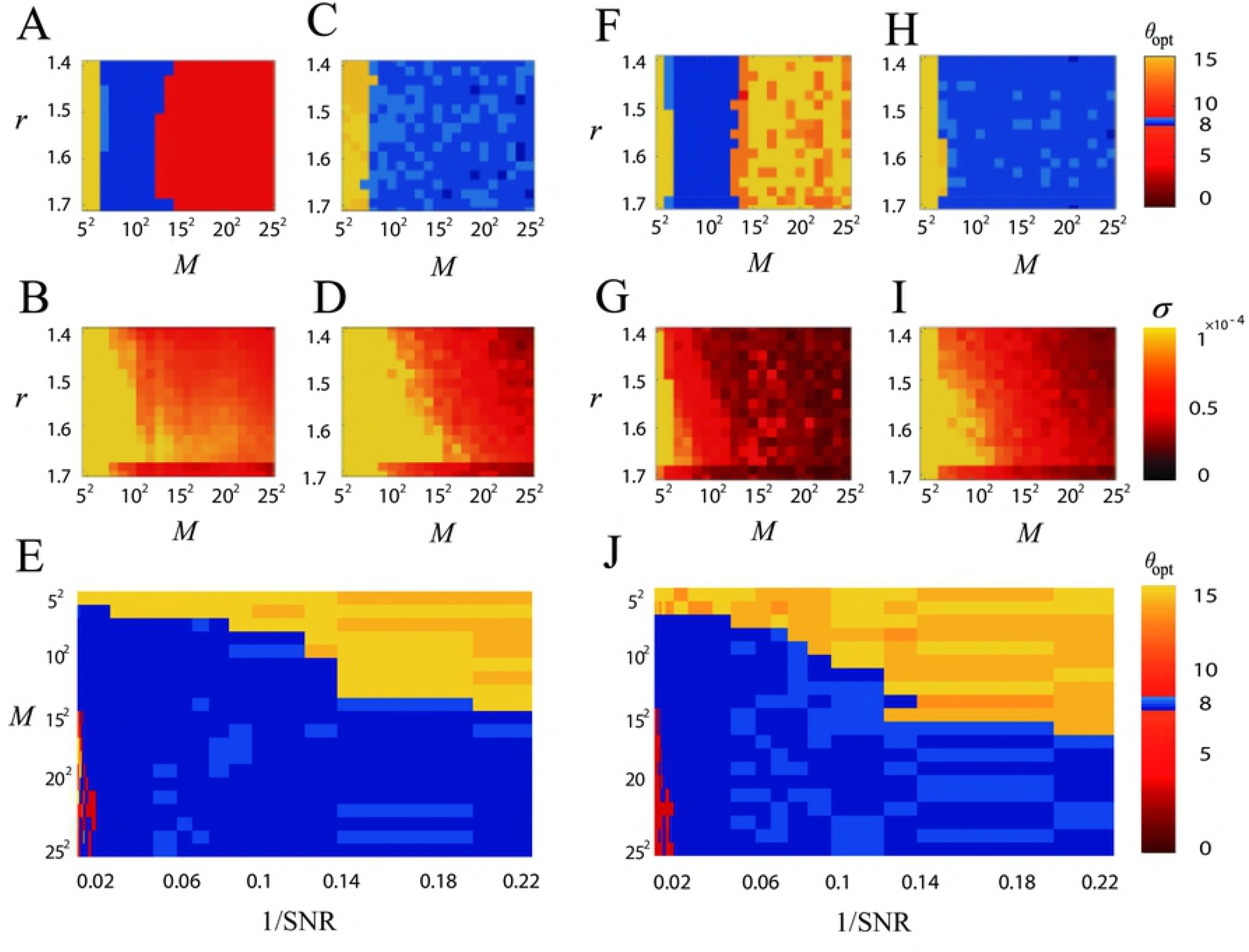
Optimal orientation associated with highest Hamming distance H. (**A**) Optimal orientation *θ*_opt_ associated with highest 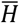 as a function of both geometrical ratio *r* an *M*, in the absence of noise. (**B**) Standard deviation *σ* of 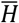 of different belts at *θ*_opt_ as a function of both *r* and *M* in the absence of noise. (**C**) *θ*_opt_ as a function of both *r* and *M* in the presence of noise. (**D**) Standard deviation of 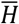 of different belts at *θ*_opt_ as a function of both *r* and *M* in the presence of noise. (**E**) *θ*_opt_ as a function of both noise 1/SNR and *M* in the presence of noise. In (**A**)-(**E**), the intensity of elliptic distortion is *ε* = 1. The color bar of (**A**) and (**C**) represents *θ*_opt_. The color bar of (**B**) and (**D**) represents standard deviation σ. In (**E**) geometrical ratio *r* = 1.5 and the environmental size is 1.5 × 1.5m^2^. (**F**)-(**J**) are the same as (**A**)-(**E**) but the intensity of elliptic distortion i**`s`** *ε* = 1.25. In (**C**) and (**H**), the effect of noise 1/SNR = 0.08. In all simulation, *f*_max_ = 200Hz.

**S5 Fig.**
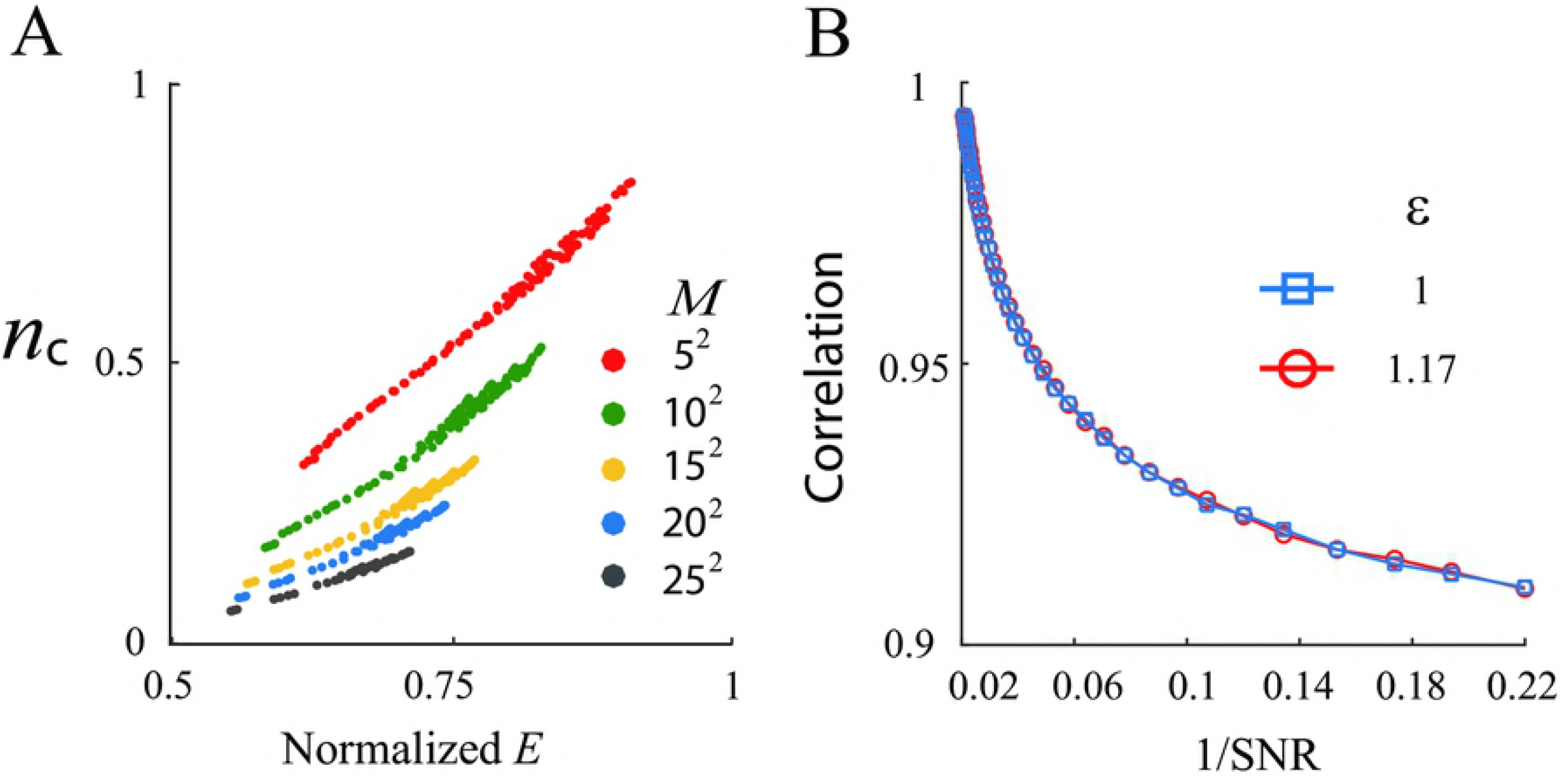
Correlation between the number of cells engaged in encoding and information entropy. (**A**) In the absence of noise, the number of cells *n*_c_ engaged in encoding belts as a function of normalized spatial information entropy *E* for different values of *M*. Normalized *E* is defined as the ratio of *E* over the maximum *E* in all belts. (**B**) Pearson correlation between *n*_c_ and *E* as a function of noise 1/SNR for two different intensities *ε* of elliptic distortion.

**S6 Fig.**
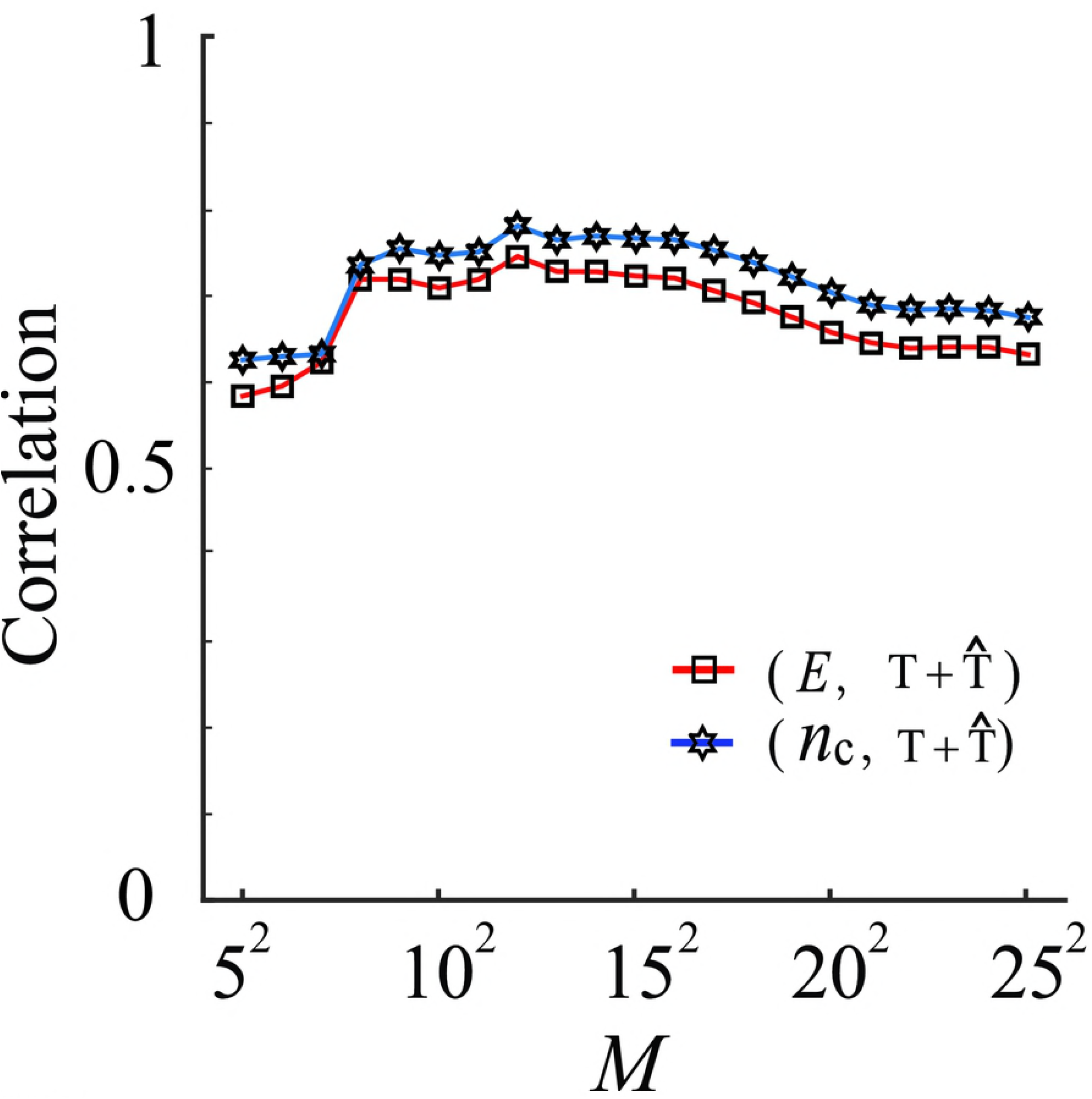
Correlation among information entropy, the number of cells engaged in encoding and the minimum repeating distance. The correlation between *E* and 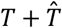 as a function of *M* and the correlation between *n*_c_ and 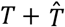 as a function of *M*. Here *M* is the number of grid cells in each module.

**S7 Fig.**
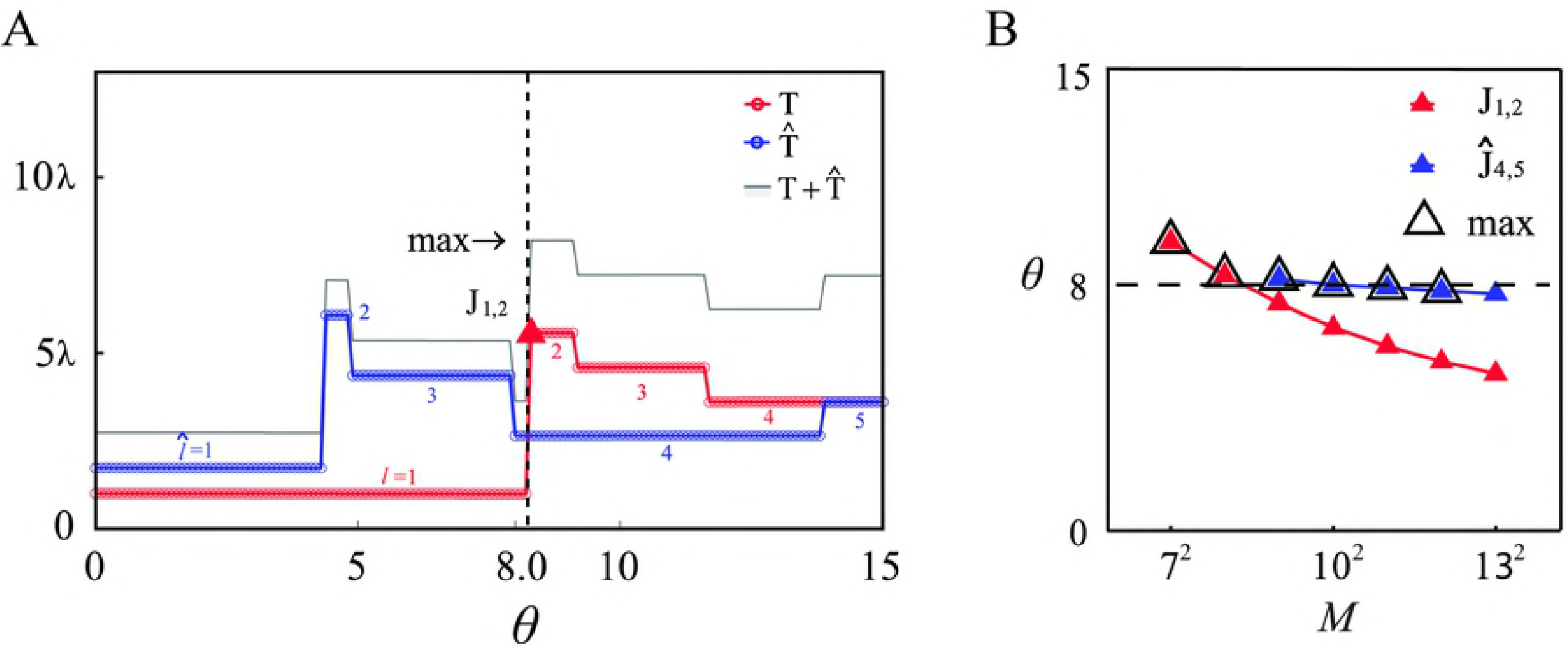
Minimum repeating distance and optimal *θ*_0_. (**A**) Simulation results of *T*, 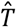 and 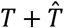 as a function of the orientation *θ* for *M* = 8^2^, which exhibit a number of abrupt transition points and plateaus. The plateaus associated with *T* and 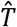 are denoted by *l* and 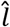, respectively, where *l* = 1,···,4 and 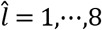, with the respective transition points as *J*_*l,l* + 1_ and 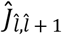 The maximum value of 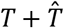 occurs at *J*_1,2_. (**B**) Simulated orientation *θ* at *J*_1,2_ and 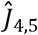 as a function of *M*. For *M* ∈ [7^2^,12^2^], *θ*_opt_ occurs at either *J*_1,2_ or 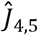, marked by the hollow triangular symbols.

**S8 Fig.**
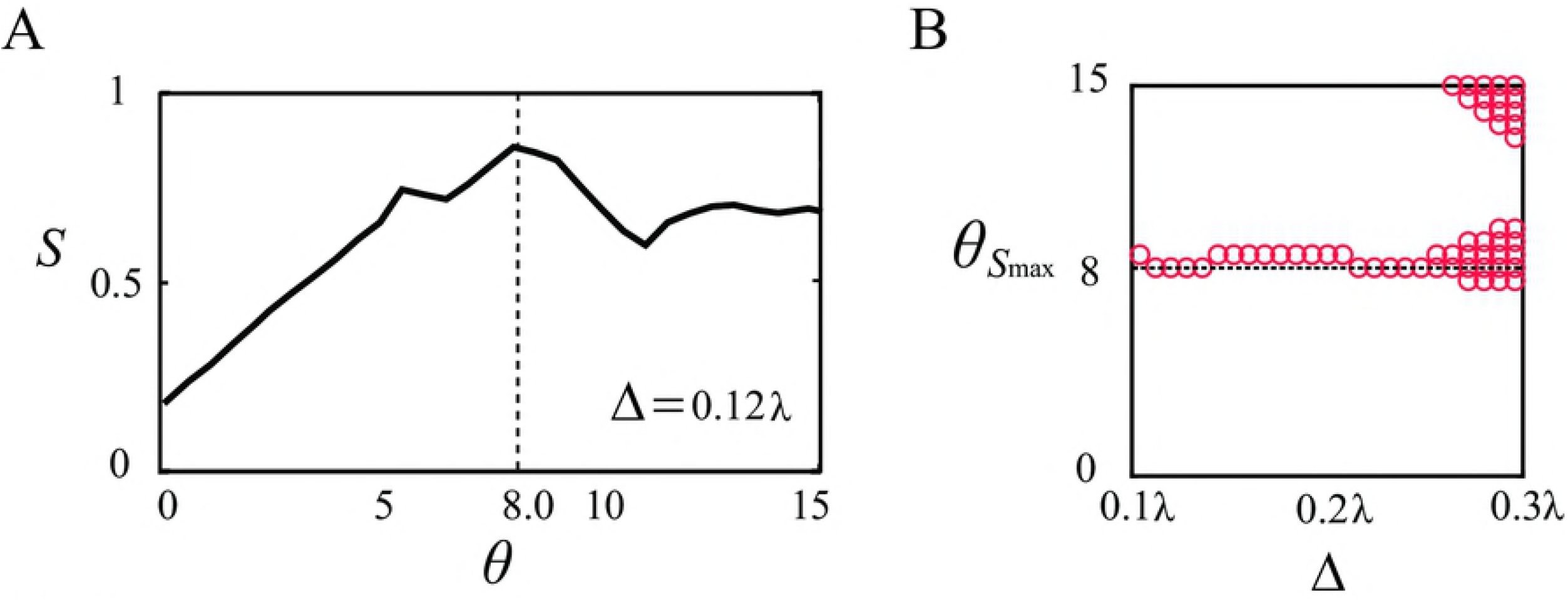
Area covered by a set of belts. (**A**) Area *S* covered by the belts as a function of *θ* for belt width Δ = 0.12λ. The largest area is achieved at *θ* = 8 degrees. (**B**) Orientation *θ*_*S*max_ associated with maximum value *θ*_*S*max_ of *S* as a function of the belt width Δ. In a wide range of Δ, 8 degrees is the exclusive optimal orientation.

**S9 Fig.**
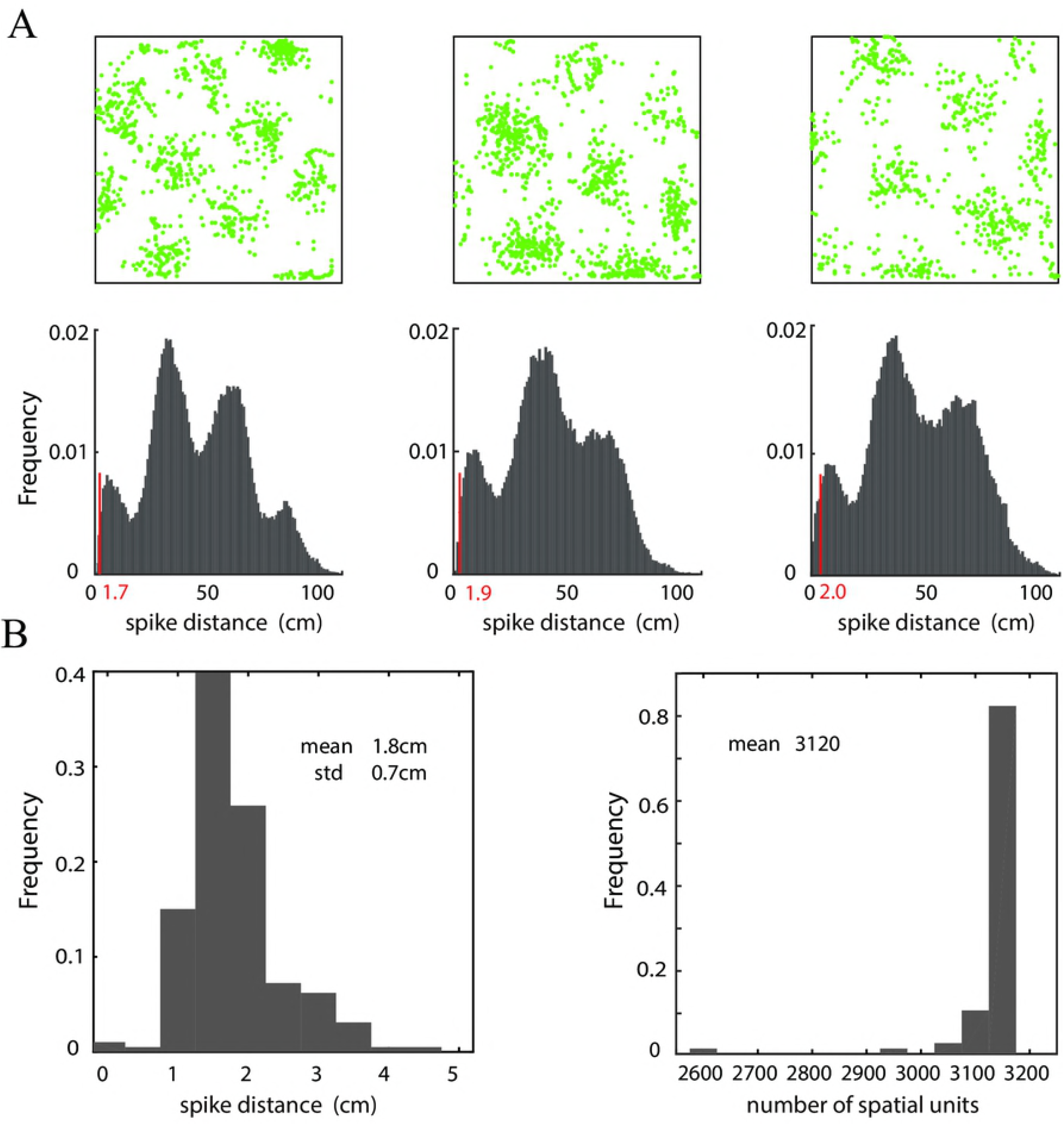
Estimation of *τ* from experiments. (**A**) Top: typical firing patterns of three grid cells, where firing locations are marked in green dots. Below: distribution of the distance between any two pairs of firing locations. The top 0.5% of smallest distance in the distribution is set to be a threshold, as marked by a red line. This threshold measures the minimal spatial resolution of rats and reflects the scale of spatial units for estimating *τ*. (B) Left: distribution of the unit scale of 193 grid cells documented in experiments. The mean distance is 1.8cm. It is set to be the average scale of the spatial unit in the environment (1.8 × 1.8_c_m^2^). Right: distribution of the numbers of spatial units computed using 76 trajectories (during about 10 minutes for each trajectory) in experiments. The mean number of spatial units containing trajectories is 3120. Thus, the average duration *τ* at every spatial unit is 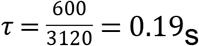.

**S10 Fig.**
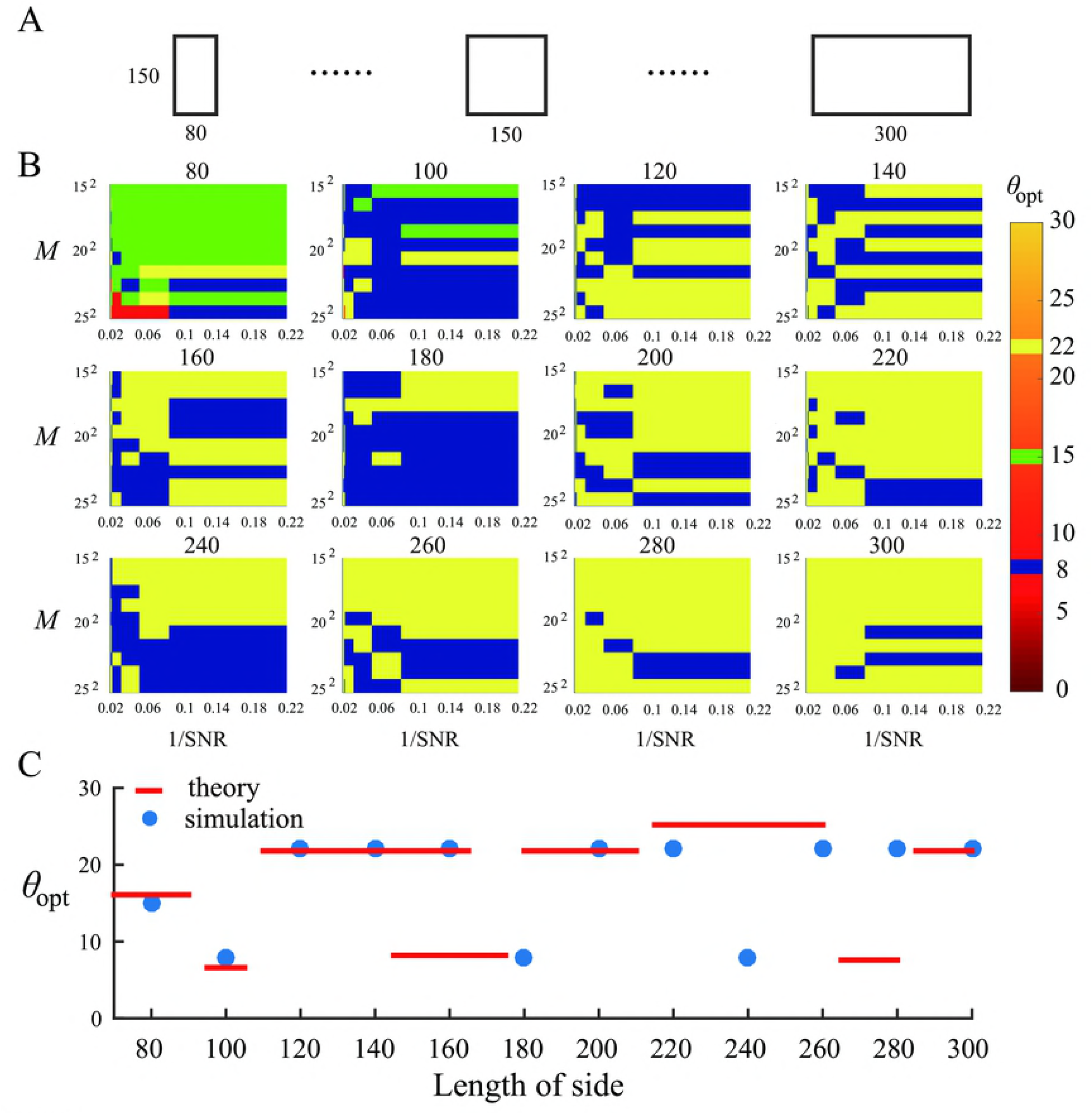
Optimal orientation in rectangular environments. (**A**) Variation of the length of a side of a rectangle. (**B**) Simulation results of *θ*_opt_ as a function of *M* and noise 1/SNR for different lengths of variant side of rectangles. The color bar represents the value of *θ*_opt_. The geometrical ratio *r* is 1.5. *θ*_opt_ is identified by the highest spatial information entropy. (**C**) Comparison between theoretical and simulation results of optimal orientation in rectangular environments. The theoretical results are obtained from Eq. (25) in the main text and the simulation results are obtained using the dominant angle for different length of the variant side in (**A**). The geometrical ratio *r* is 1.5 and *f*_max_ = 200Hz.

